# A Major Role of Class III HD-ZIPs in Promoting Sugar Beet Cyst Nematode Syncytium Formation in *Arabidopsis*

**DOI:** 10.1101/2024.02.15.580549

**Authors:** Xunliang Liu, Melissa G. Mitchum

**Affiliations:** Department of Plant Pathology and Institute of Plant Breeding, Genetics, and Genomics, University of Georgia

**Keywords:** Beet Cyst Nematode, CLAVATA, CLE, Peptide Hormone, Effector, HD-ZIP III, RNA-seq, Stem Cell Organizer, Syncytium

## Abstract

Cyst nematodes use a stylet to secrete plant CLE-like peptide effector mimics into selected root cells of their host plants to hijack CLE signaling pathways for feeding site (syncytium) formation. Here, we identified *ATHB8*, a HD-ZIP III family transcription factor, as a downstream component of the CLE signaling pathway in syncytium formation. *ATHB8* is expressed in the early stages of syncytium initiation, and then transitions to neighboring cells of the syncytium as it expands; an expression pattern coincident with auxin response at the infection site. Conversely, *MIR165a*, which expresses in endodermal cells and moves into the vasculature to suppress HD-ZIP III TFs, is down-regulated near the infection site. Knocking down HD-ZIP III TFs by inducible over-expression of *MIR165a* in *Arabidopsis* dramatically reduced female development of the sugar beet cyst nematode (*Heterodera schachtii*). HD-ZIP III TFs are known to function downstream of auxin to promote cellular quiescence and define stem cell organizer cells in vascular patterning. Taken together, our results suggest that HD-ZIP III TFs may function as a connecting point for CLE and auxin signaling pathways in promoting syncytium formation, possibly by inducing root cells into a quiescent status and priming them for initial syncytial cell establishment and/or subsequent cellular incorporation.

## Introduction

Obligate sedentary endoparasitic cyst nematodes (CNs) require a reliable and sustained local source of nutrition within host roots to complete their life cycle. They achieve this by establishing a unique and highly specialized feeding site called a syncytium in host roots. The syncytium forms when an infective juvenile selects a single root cell, usually a procambium or pericycle cell (Golinowski et al., 1996; Holtmann et al., 2000; Sobczak and Golinowski, 2009; Anjam et al., 2020), in the vasculature of a host root, and secretes a suite of effectors through a stylet to induce massive re-programming of the selected host cell (Hewezi and Baum, 2013; Molloy et al., 2023). The syncytium expands along the vascular cylinder by gradually incorporating neighboring cells by partial cell wall dissolution and protoplasmic fusion (Gipson et al., 1971; Golinowski et al., 1996; Grundler et al., 1998; Ohtsu et al., 2017). Researchers have strived to understand the signaling events regulating the highly orchestrated process of syncytium formation. These efforts have shed light on the critical role of CN effector-mediated modulation of phytohormone and peptide hormone signaling pathways in syncytium formation, most notably through the modulation of auxin and CLE signaling pathways (Gheysen and Mitchum, 2019).

At the early stage of infection, perturbations to auxin homeostasis are key to syncytium establishment. A local accumulation of auxin is pivotal for syncytium initiation and is associated with changes in auxin biosynthesis, polar transport, and signaling (Grunewald et al., 2009; Lee et al., 2011; Habash et al., 2017; Oosterbeek et al., 2021; Suzuki et al., 2022). Several CN effectors have been identified to directly modulate auxin accumulation and signaling at the infection site. The beet cyst nematode (BCN) tyrosinase-like effector increases auxin content and promotes BCN susceptibility when ectopically expressed in *Arabidopsis* (Habash et al., 2017); the BCN effector 19C07 directly interacts with *Arabidopsis* Like-AUX1 3 (LAX3) auxin influx protein and is likely to increase LAX3 activity and promote auxin flow into the syncytium and adjacent cells (Lee et al., 2011); while the BCN effector 10A07 is translocated into the plant cell nucleus after being phosphorylated by plant protein kinase IPK (AT2G37840) and interacts with Indoleacetic Acid-Induced Protein 16 (IAA16), a suppressor of Auxin Responsive Factors (ARFs) to possibly release corresponding ARFs for auxin signaling (Hewezi et al., 2015). Host genes related to auxin biosynthesis, distribution, and signaling are also modulated by CN at the infection site. *YUCCA4* (*YUC4*), encoding a member of YUCCA family enzymes that mediates a speed limiting step in the main auxin biosynthesis pathway, is up-regulated at the CN infection site, along with a few other auxin biosynthesis enzymes (Guarneri et al., 2022; Suzuki et al., 2022). Auxin efflux *PIN1* and *PIN7* genes are down-regulated at the early feeding site to prevent out flow of auxin from the initial syncytial cell, while PIN3 and PIN4 proteins are re-localized to the lateral membranes of the expanding syncytium to re-distribute auxin for lateral expansion of the syncytium (Grunewald et al., 2009). Of the 22 *ARF* genes in *Arabidopsis*, all except one (*ARF8*) were found to be up-regulated at the infection site, with distinct and overlapping spatiotemporal expression patterns, indicating *ARFs* may play different roles in various stages of syncytium formation (Hewezi et al., 2014). At the same time, *miRNAs* target *ARF* mRNAs for degradation, and several *IAA* genes, which encode suppressors of ARF activities, are down-regulated at the infection site (Ithal et al., 2007; Hewezi et al., 2008a), further highlighting the sophisticated manipulation of auxin signaling for feeding site formation.

CNs also co-opt developmental programs of host cells through deployment of CLAVATA3 (CLV3)/ENDOSPERM SURROUNDING REGION (ESR) (CLE) peptide effectors (Wang et al., 2001; Wang et al., 2005; Lu et al., 2009; Wang et al., 2011; Guo et al., 2017; Mitchum and Liu, 2022). In plants, CLEs are expressed as prepropeptides, and are then secreted into the apoplast where they are processed into mature 12-14 amino acid CLE peptides (Ito, 2006; Ni and Clark, 2006; Ohyama et al., 2008; Ni et al., 2011). CLE peptides are perceived on the cell surface by leucine-rich repeat receptor-like kinases (LRR-RLKs) and generally function in meristem cell homeostasis (Dropkin, 1969; Hirakawa et al., 2008; Ogawa et al., 2008; Morita et al., 2016). In nematodes, CLE effectors are injected through a stylet into the host cell cytoplasm in the form of a proprotein (Wang et al., 2010b), and are then re-secreted through the endomembrane system into the apoplast (Wang et al., 2010a; Wang et al., 2010b; Wang et al., 2021), where they are processed into mature CLE peptides (Ni and Clark, 2006; Guo et al., 2011; Ni et al., 2011). This re-secretion process not only allows CLE effectors to gain host-specific modifications that are critical for CLE activity (Ito, 2006; Ohyama et al., 2008; Ohyama et al., 2009; Guo et al., 2011; Chen et al., 2015), but it also gives the nematode access to the CLE signaling pathways of adjacent cells to prime these cells for syncytium incorporation (Wang et al., 2010b; Wang et al., 2021; Mitchum and Liu, 2022).

Plant CLEs can be classified into A- and B-types, based on their ability to promote terminal differentiation of shoot and root apical meristem cells (Whitford et al., 2008). Both A- and B-type CLEs function in a CLE-Receptor-Like Kinase (RLK)-WUSCHEL(WUS)/WUSCHEL-LIKE HOMEOBOX(WOX) paradigm to regulate stem cell activities. While A-type CLEs mainly functions in restricting plant shoot and root apical meristems (SAM/RAM) (Fletcher, 2020), B-type CLE, namely TRACHEARY ELEMENT DIFFERENTIATION INHIBITORY FACTOR (TDIF) (Ito, 2006), signals through TDIF receptor (TDR) to regulate a feed forward network to promote vascular (pro)cambial cell proliferation (Etchells and Turner, 2010; Hirakawa et al., 2010; Ji et al., 2010; Etchells et al., 2013; Etchells et al., 2016; Smit et al., 2020). Interestingly, B-type CLE promotion of vascular stem cell proliferation can be enhanced by co-treatment of A-type CLEs like CLE6p, CLV3p, and CLE19p (Whitford et al., 2008).

Although the molecular mechanism of synergistic action of A- and B-type CLEs is not clear, some *Arabidopsis* endogenous A-type CLEs do express in vascular tissue (Jun et al., 2010), and some showed activity in regulating vascular cell differentiation (Hirakawa et al., 2011; Kondo et al., 2011; Depuydt et al., 2013; Rodriguez-Villalon et al., 2014; Qian et al., 2018; Hu et al., 2022; Carbonnel et al., 2023). Remarkably, molecular mimics of both plant A- and B-type CLE peptides are found in CNs (Wang et al., 2001; Wang et al., 2005; Lu et al., 2009; Wang et al., 2011; Guo et al., 2017; Mitchum and Liu, 2022). Both A- and B-type nematode CLE effectors signal through their respective receptors to promote cyst nematode parasitism (Guo et al., 2011; Replogle et al., 2011; Replogle et al., 2012; Guo et al., 2015; Guo et al., 2017), by, at least partially, activating *WOX4* gene expression (Guo et al., 2017).

Another important factor in regulating vascular stem cell homeostasis is the class III HOMEODOMAIN LEU-ZIPPER (HD-ZIP III) family of transcriptional factors (Ramachandran et al., 2017). The *Arabidopsis* HD-ZIP III family has five members, including *ATHB8*, *ATHB15/CORONA* (*CNA*), *PHABULOSA* (*PHB*), *PHAVOLUTA* (*PHV)*, and *REVOLUTA* (*REV*). Expression of HD-ZIP IIIs is highly regulated by auxin biosynthesis and polar auxin transport (Bishopp et al., 2011; Ursache et al., 2014), and at least one member, *ATHB8*, is directly regulated by AUXIN RESPONSE FACTOR 5 (ARF5)/MONOPTEROS (MP) (Donner et al., 2009). In the vasculature, HD-ZIP III genes show peak expression on the xylem side of vascular cambium (Smetana et al., 2019), correlating with peak auxin concentrations (Uggla et al., 1996; Immanen et al., 2016). Post-transcriptionally, HD-ZIP III levels are regulated by *microRNA 165/166* (Emery et al., 2003; Mallory et al., 2004; Kim et al., 2005; Williams et al., 2005; Zhong and Ye, 2007; Carlsbecker et al., 2010; Miyashima et al., 2011). *MIR165/166* are activated in the endodermis, by stele-derived SHORT-ROOT (SHR) (Helariutta et al., 2000; Gallagher et al., 2004; Carlsbecker et al., 2010), and then diffuse back into the stele to establish a gradient of HD-ZIP III levels, where high HD-ZIP III level specifies metaxylem and low HD-ZIP III level specifies protoxylem (Carlsbecker et al., 2010; Miyashima et al., 2011). Recently, Smetana *et al* (2019) established that the xylem-identity cell, which is promoted by high auxin and HD-ZIP III genes, functions as a stem cell organizer (SCO) to promote adjacent stem cell division and overall vascular patterning, reminiscent of the OC in SAM and QC in RAM (Smetana et al., 2019).

Crosstalk between CLE and auxin signaling pathways regulates stem cell homeostasis and vascular patterning (Ji et al., 2010; DiGennaro et al., 2018; Han et al., 2018; Jones et al., 2021; John et al., 2023), and both CLE and auxin are known to be involved in CN-induced syncytium formation (Gheysen and Mitchum, 2019). Nevertheless, it is not clear how these two phytohormones work together to promote syncytium formation. Here, we identified *ATHB8*, an HD-ZIP III family transcription factor, as a downstream factor of the HsCLE2 peptide effector, providing a potential intersection point for CLE and auxin signaling pathways in syncytium formation. We also demonstrated that *ATHB8*, along with other HD-ZIP III family genes, plays an important role in syncytium formation during beet cyst nematode infection in *Arabidopsis*. Distribution of *ATHB8* expression suggests that HD-ZIP III genes may function in both initial syncytial cell establishment and incorporation of adjacent cells for syncytium expansion, by promoting quiescence status of root cells.

## Results

### Gene Expression Profiles of the *clv1-101 clv2-101 rpk2-5* and Wild Type Roots Converge Upon BCN Infection

Our lab has previously reported that *Arabidopsis* receptor-like kinases CLV1, CLV2, and RPK2 perceive *Heterodera schachtii* (*H. schachtii*, BCN) CLE-like peptide effectors to promote parasitism (Replogle et al., 2011; Replogle et al., 2012), and that *WOX4* functions downstream of these CLE receptors to promote syncytium formation (Guo et al., 2017). To isolate additional downstream CLE peptide signaling components involved in BCN parasitism, we profiled the transcriptome of BCN infection sites from both wild-type and *clv1-101 clv2-101 rpk2-5* triple (hereafter *clv* triple) CLE receptor mutant plants (Replogle et al., 2012; Guo et al., 2017). To filter out genes specifically responsive to nematode CLE effectors, we also included a group of samples treated with synthetic HsCLE2p effector (Figure 1A), a mimic of *Arabidopsis* CLE5/6p (Wang et al., 2011). The treatment dose and timepoint was optimized to 5 μM HsCLE2p for 3 hours, using the expression of *WOX4* as the indicator (Supplemental Figure S1A) (Guo et al., 2017). The collected samples were verified for *WOX4* expression before proceeding to library construction (Supplemental Figure S1B).

**Figure 1.**
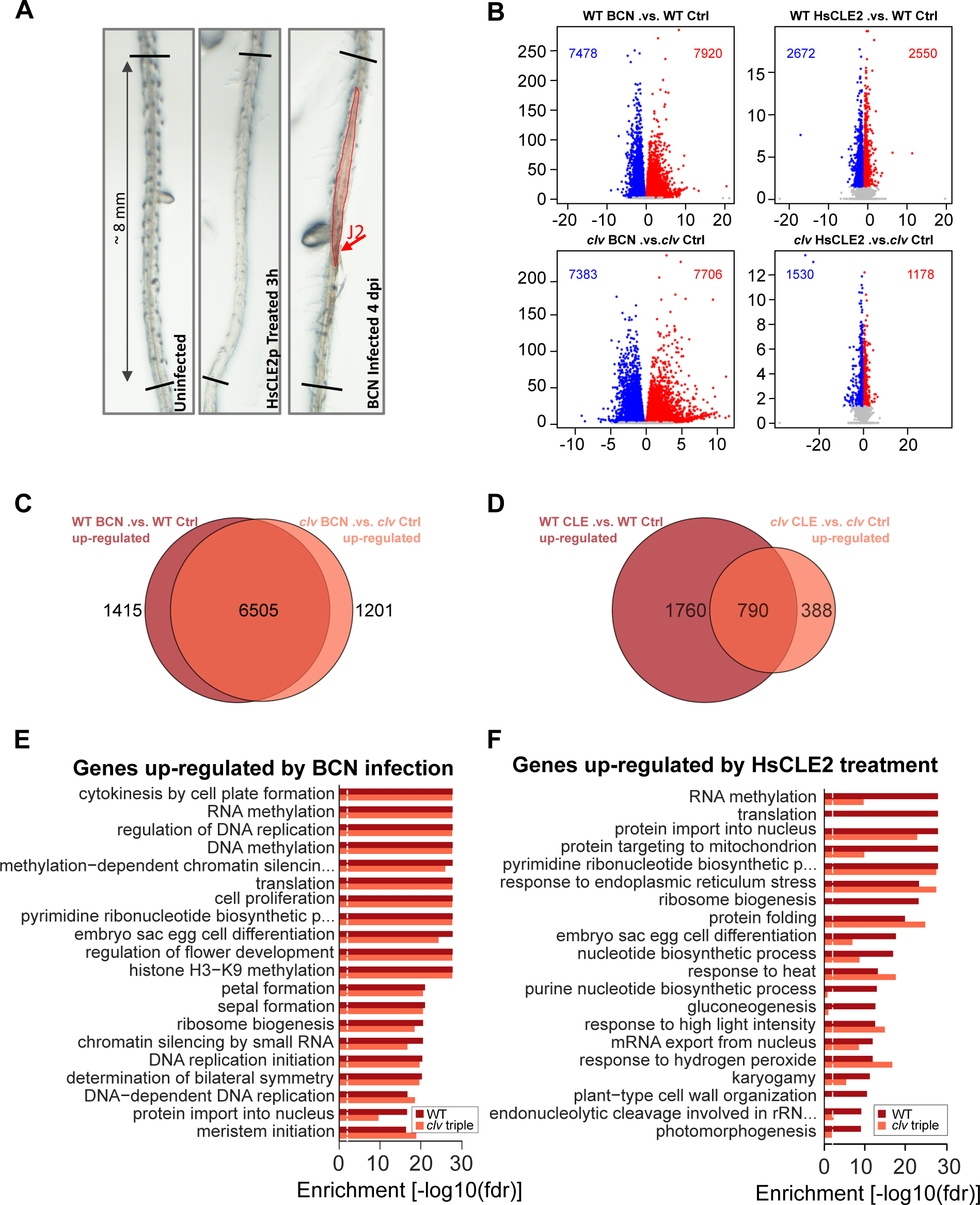
Transcriptome profiling of BCN infection sites in wild-type and the clv1-101 clv2-101 rpk2-5 triple CLE receptor mutant. **A.** Example of excised root segments used for RNA sequencing. Approximately 8 mm long root segments spanning BCN infection sites and corresponding uninfected or HsCLE2p-treated root tissues were cut under a stereoscope and immediately frozen in liquid nitrogen for RNA isolation. **B.** Volcano plot of differentially expressed genes upon BCN infection and HsCLE2p treatment in wild-type and the *clv* triple mutant. **C.** Venn diagram showing overlap of up-regulated genes in wild-type and *clv* triple mutant upon BCN infection. **D.** Venn diagram showing overlap of up-regulated genes in wild-type and *clv* triple mutant upon HsCLE peptide treatment. **E.** BCN up-regulated genes in wild-type and *clv* triple mutant were enriched in similar GO terms of biological process. **F.** HsCLE2p up-regulated genes in wild-type and *clv* triple mutant were enriched in different GO terms for biological process.

RNA sequencing yielded over 25 million reads from each sample (Supplemental Table S1). All marker genes were expressed as expected (Supplemental Figure S2A). *clv1-101* and *clv2-101* are null alleles and *CLV1* and *CLV2* expression was barely detectable in the *clv* triple mutant; whereas *RPK2* showed high expression in the mutant due to *rpk2-5* allele being a point mutation (Supplemental Figure S2D; S. Sawa, personal communication). Expression of *WOX4* and *CLE41*, both known to function downstream of CLE signaling in BCN infection (Guo et al., 2017), were induced by HsCLE2p treatment in wild-type samples but not in the *clv* triple mutant (Supplemental Figure S2A), indicating effective HsCLE2p treatment. PCA analysis showed that samples clustered as expected (Supplemental Figure S2B), except that Rep 1 of HsCLE2p treated *clv* triple mutant clustered closer to wild-type samples rather than *clv* triple mutant samples (Supplemental Figure S2B). Similarly, this sample also clustered with HsCLE2 treated wild-type samples in hierarchical clustering (Supplemental Figure S2C). Despite the expression of all marker genes in this sample resembling the other two biological replicates of HsCLE2p treated *clv* triple mutant (Supplemental Figure S2A), and confirmation of the *rpk2-5* mutation in this sample (Supplemental Figure S2D), the sample was removed from the downstream analysis out of caution.

Differentially expressed genes (DEGs) were identified with the DEseq2 package (Love et al., 2014). In wild-type roots, 7,920 and 7,478 genes were up- or down-regulated, respectively, by BCN infection (padj < 0.05) (Figure 1B, Supplemental Dataset S2), and 2,550 and 2,672 genes were up- or down-regulated by HsCLE2p treatment (Figure 1B, Supplemental Dataset S2). In *clv* triple mutant roots, 7,706 and 7,383 genes were up- or down-regulated by BCN infection, and 1,178 and 1,530 genes were up- or down-regulated by HsCLE2 treatment (Figure 1B, Supplemental Dataset S2). Of the 7,920 up- and 7,478 down-regulated genes by BCN infection in the wild-type, 6,505 (82.1%) and 6,337 (84.7%) genes were also up- or down-regulated, respectively, by BCN infection in the *clv* triple mutant (Figure 1C; Supplemental Figure S3A). In comparison, with HsCLE2p treatment, only 46.2% (1,178 out of 2,550) of up-regulated and 36.5% (976 out of 2,672) of down-regulated genes showed the same regulatory trend between the *clv* triple mutant and the wild-type (Figure 1D; Supplemental Figure S3B). The higher percentage overlap of BCN-induced DEGs between the *clv* triple mutant and wild-type was also confirmed by Gene Ontology enrichment. The top 20 most enriched Biological Processes in BCN up- or down-regulated genes in wild type, were also enriched in the BCN up- or down-regulated DEGs in the *clv* triple mutant, often to a comparable enrichment level (Figure 1E, Supplemental Figure S3C, Supplemental Dataset S3). Whereas for HsCLE2p treatment, enriched GO terms in the *clv* triple mutant showed greater differences from that of the wild-type (Figure 1F, Supplemental Figure S3D, Supplemental Dataset S3). These results suggested that BCN infection triggered a similar gene expression response in the *clv* triple mutant compared to that of the wild-type, despite the lack of CLV1, CLV2, and RPK2 receptors. Interestingly, far less DEGs were identified between BCN-infected mutant and wild-type samples (Supplemental Dataset S2, *clv* BCN vs. WT_BCN, 2252 DEGs) compared to the number of DEGs found between uninfected *clv* triple mutant and wild type samples (Supplemental Dataset S2, *clv*_Ctrl .vs. WT_Ctrl, 5,388 DEGs). In contrast, HsCLE2p treatment resulted in an increased number of DEGs between mutant and wild-type (Supplemental Dataset S2, *clv*_HsCLE2 vs. WT_HsCLE2, 6440 DEGs) compared to untreated samples. Of the 5,388 DEGs between the *clv* triple mutant and wild-type, only 874 were differentially expressed following BCN infection (Supplemental Figure S4A, B; Supplemental Dataset S2) compared to 3,736 following HsCLE2p treatment (Supplemental Figure S4C, D). Again, these results suggested that BCN infection reduced gene expression differences between the transcriptome of the *clv* triple mutant and that of the wild-type. Consistently, BCN-infected wild-type and mutant samples clustered closer to each other compared to uninfected wild-type and mutant samples (Supplemental Figure S2B, C), indicating increased similarity between these two genetic backgrounds following nematode infection. The observed transcriptomic similarity between the *clv* triple mutant and wild-type samples in response to BCN infection indicated that BCN infected samples might not be the optimal choice for filtering out downstream genes of CLE signaling. Instead, HsCLE2p treatment would likely be a more effective approach for this purpose.

### Identification of Downstream CLE Signaling Genes Involved in BCN Infection

We reasoned that genes promoted by CLE effectors via CLE receptors should have reduced expression levels in the mutant, due to attenuated endogenous CLE signaling, and should have an attenuated response to HsCLE2p treatment in the *clv* triple mutant. Moreover, these genes should also be promoted by BCN infection. Thus, we filtered genes that were, 1) down-regulated in the *clv* mutant compared to wild-type when uninfected, 2) up-regulated by HsCLE2p treatment in the wild-type root but not in the *clv* triple mutant, and 3) up-regulated by BCN infection. A total of 147 genes fit these criteria (Figure 2A-B; Supplemental Dataset S4). Using a similar approach, a set of 176 genes was identified as potential targets suppressed by the HsCLE2p effector (Supplemental Figure S7; Supplemental Dataset S5). For the time being, we primarily focused on the 147 HsCLE2p-promoted candidate genes. These 147 candidate genes were further divided into three tiers based on the level of induction in response to BCN infection, and their annotated molecular function (Supplemental Dataset S4). Genes having a log2 fold change larger than 2, or with regulatory related functions, were categorized as tier 1. To verify the expression pattern of these genes with qPCR, a new set of reference genes was selected and tested (Supplemental Figure S5B - C), since previously published reference genes showed expression variations among our samples (Supplemental Figure S5A). Thirty-two of the 38 tier 1 genes were selected for qPCR verification, of which most followed the expression pattern shown in the RNA sequencing data with few exceptions (Supplemental Figure S6).

**Figure 2.**
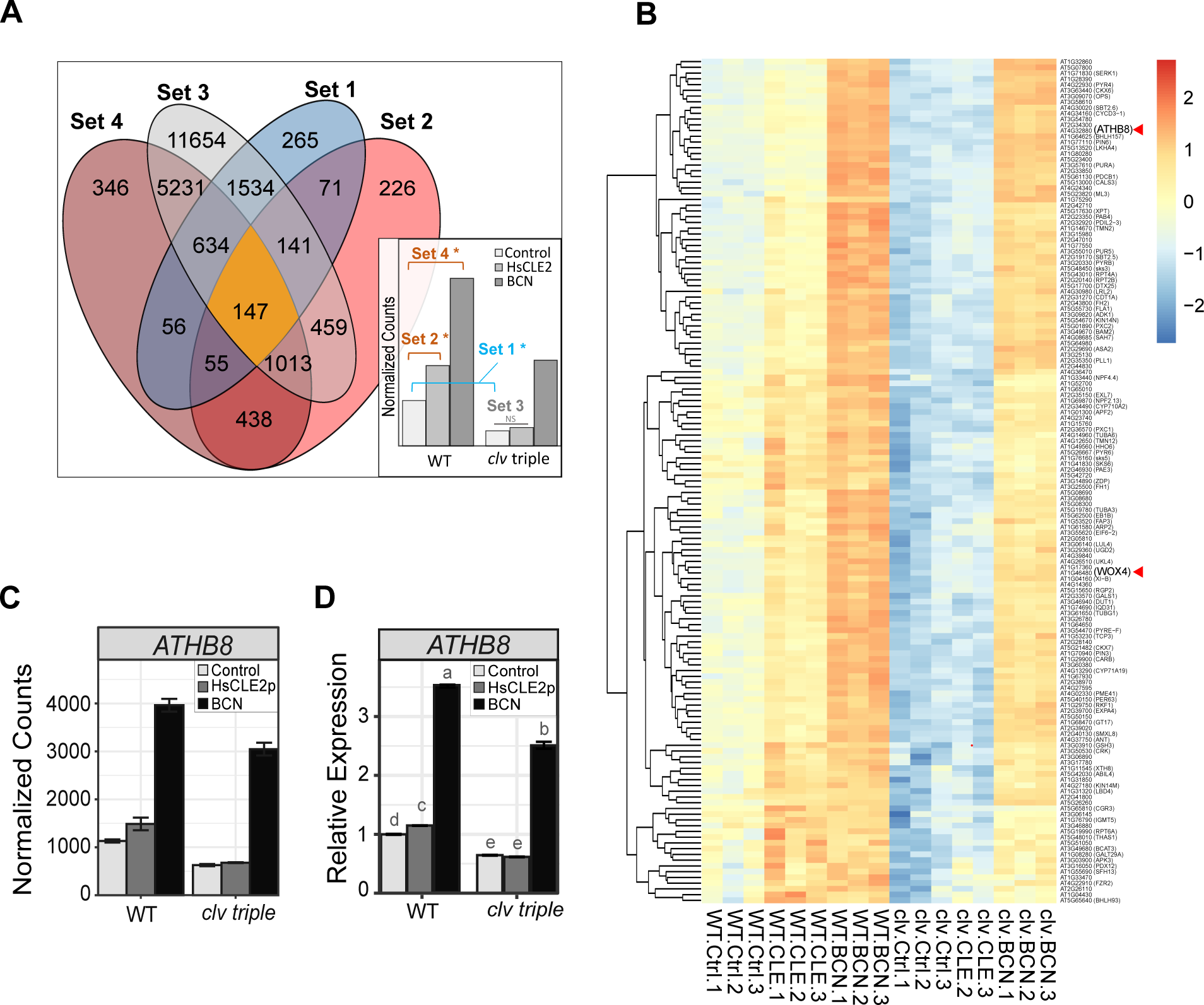
Identification of potential genes downstream of CLE signaling upon BCN infection. **A.** Criteria used for filtering CLE downstream genes. Genes (147) that fit these four criteria were considered as CLE downstream genes, 1). Down-regulated in the *clv* triple mutant compared to wild-type in control sample; 2). Up-regulated by HsCLE2p treatment in wild-type seedlings, but not in the *clv* triple mutant (3); 4). Up-regulated by BCN infection in wild-type seedlings. **B.** Heatmap showing expression profiles of the 147 candidate genes positively regulated by CLE signaling upon BCN infection. Expression of *ATHB8* and *WOX4* are denoted with red arrowheads. **C.** Expression of *ATHB8* gene in the RNAseq dataset. **D.** qPCR validation of *ATHB8* gene expression in wild-type and *clv* triple mutant upon HsCLE2p treatment and BCN infection. Letter above each bar represents statistical groups of Turkey’s HSD test following ANOVA analysis.

### *ATHB8* is Up-Regulated at the Periphery of the Developing Syncytium

One tier 1 gene, *ATHB8*, in particular, was identified as a strong candidate downstream CLE signaling gene involved in BCN infection. In the *clv* triple mutant, its response to HsCLE2p treatment was completely abolished (Figure 2C). Although it was still up-regulated by BCN infection in the mutant, the expression level did not reach that of wild-type (Figure 2C), indicating impaired CLE signaling compromised *ATHB8* activation upon BCN infection. This expression pattern was also verified by qPCR (Figure 2D). *ATHB8* belongs to the HD-ZIP III transcription factor family. Members of this five-member gene family in *Arabidopsis* are well known for their roles in vascular cell fate determination and vascular patterning (Baima et al., 2001; Carlsbecker et al., 2010; Smetana et al., 2019). In addition, all five members of the HD-ZIP III family were up-regulated by BCN infection (Supplemental Figure S8), indicating that these genes were likely to be involved in the differentiation of syncytial cells.

To explore the role of *ATHB8* in nematode parasitism, we first examined the activity of the *ATHB8* promoter in response to BCN infection using a *ProATHB8::GUS* transgene. In uninfected roots, the *ATHB8* promoter is only active in the vasculature and root apical meristem (RAM). It is highly expressed in RAM and gradually decreases through the division zone. In mature roots, the *ATHB8* promoter activity becomes slightly elevated again (Supplemental Figure S9A), which is probably associated with its role in secondary vascular development (Baima et al., 2001; Smetana et al., 2019). Upon BCN infection, *ProATHB8::GUS* expression increased at the infection site compared to adjacent root tissue at early nematode developmental stages (Figure 3A). By the late J2 (2^nd^ stage Juvenile) stage and on-wards, *ATHB8* was expressed at the periphery of the syncytium, rather than in the syncytium itself (Figure 3B-E). However, *ATHB8* up-regulation become less prominent by the late J2 or J3 stage compared to the early J2 stage (Figure 3A-C). Once the nematode reached the adult female stage, *ATHB8* expression at the periphery of the syncytium was barely detectable (Figure 3D). Similarly, in male-associated syncytia, *ProATHB8::GUS* was also expressed in the periphery of syncytium but not in the syncytium itself (Figure 3E). We next examined a more detailed *ATHB8* expression at the early stage of syncytium formation using the *ProATHB8::4xYFP* transgene. In uninfected seedlings, *ProATHB8::4xYFP* showed a similar expression pattern to that of *ProATHB8::GUS* (Supplemental Figure S10A). An optical section near the root tip showed that *ATHB8* is expressed in the xylem axis in early vascular development (Supplemental Figure S10A), consistent with previous reports (Carlsbecker et al., 2010; Bishopp et al., 2011). In mature roots, *ATHB8* expression is predominantly located on the xylem side of vascular cambium (Smetana et al., 2019). Upon nematode infection, *ATHB8* expression level is clearly increased one day-post-inoculation (dpi) at the infection site, compared to adjacent root (Figure 4A). The YFP signal continues to increase up to about 4 dpi before it starts to decrease (Figure 4A). An optical section of a five-day old syncytium showed that the YFP signal is located at the periphery of the syncytium (Supplemental Figure S10B), consistent with what was observed with the *ProATHB8::GUS* transgene (Figure 3B).

**Figure 3.**
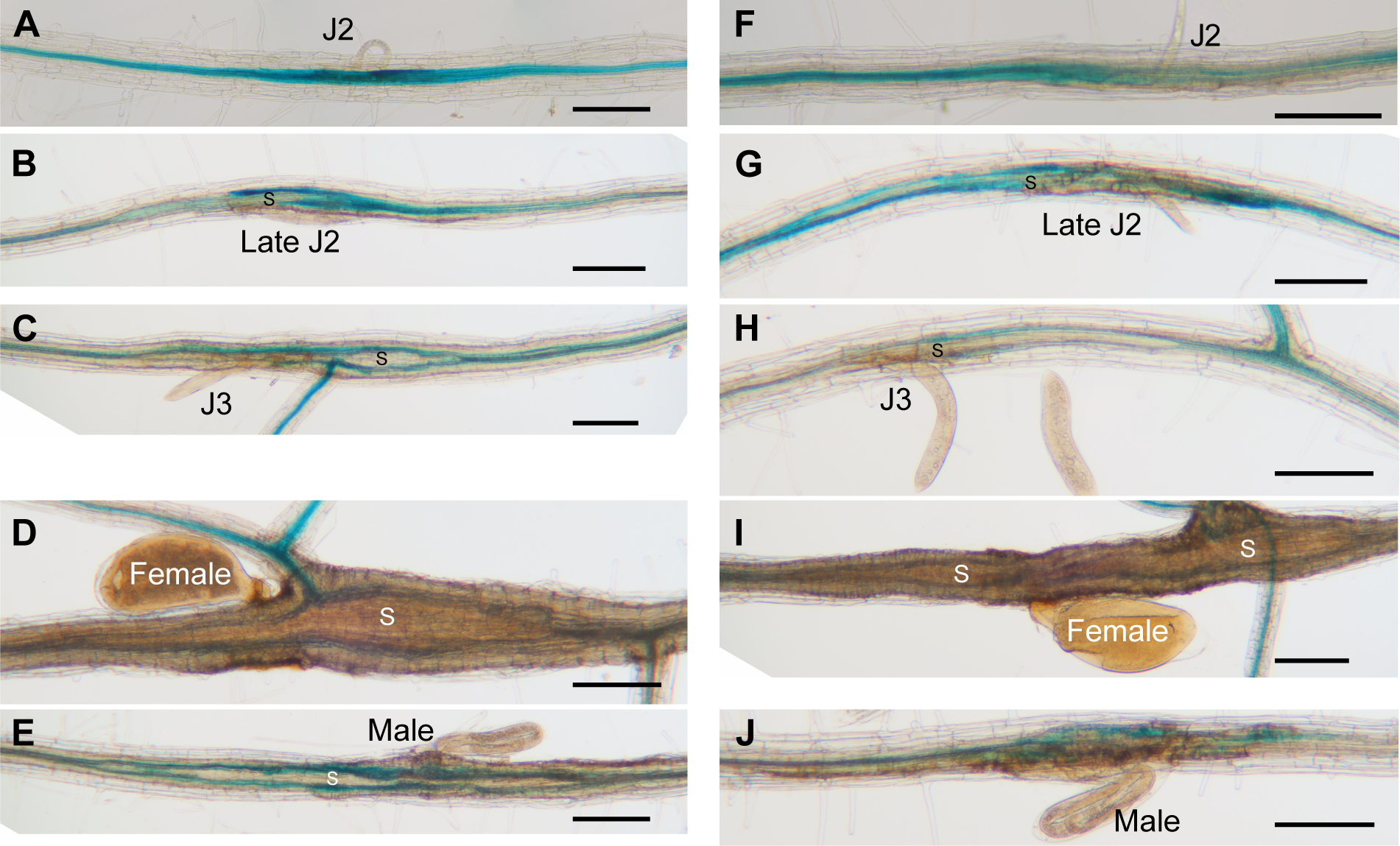
Expression of *ProATHB8::GUS* in the BCN infection site in wild-type and *clv* triple mutant. *ProATHB8::GUS* expression at different stage of syncytium development in wild-type [Ws] (**A-E**) and the *clv1-101 clv2-101 rpk2-5* triple mutant (**F-J**). J2, second-stage juvenile; J3, third-stage juvenile; S, syncytium. bar = 200 um.

**Figure 4.**
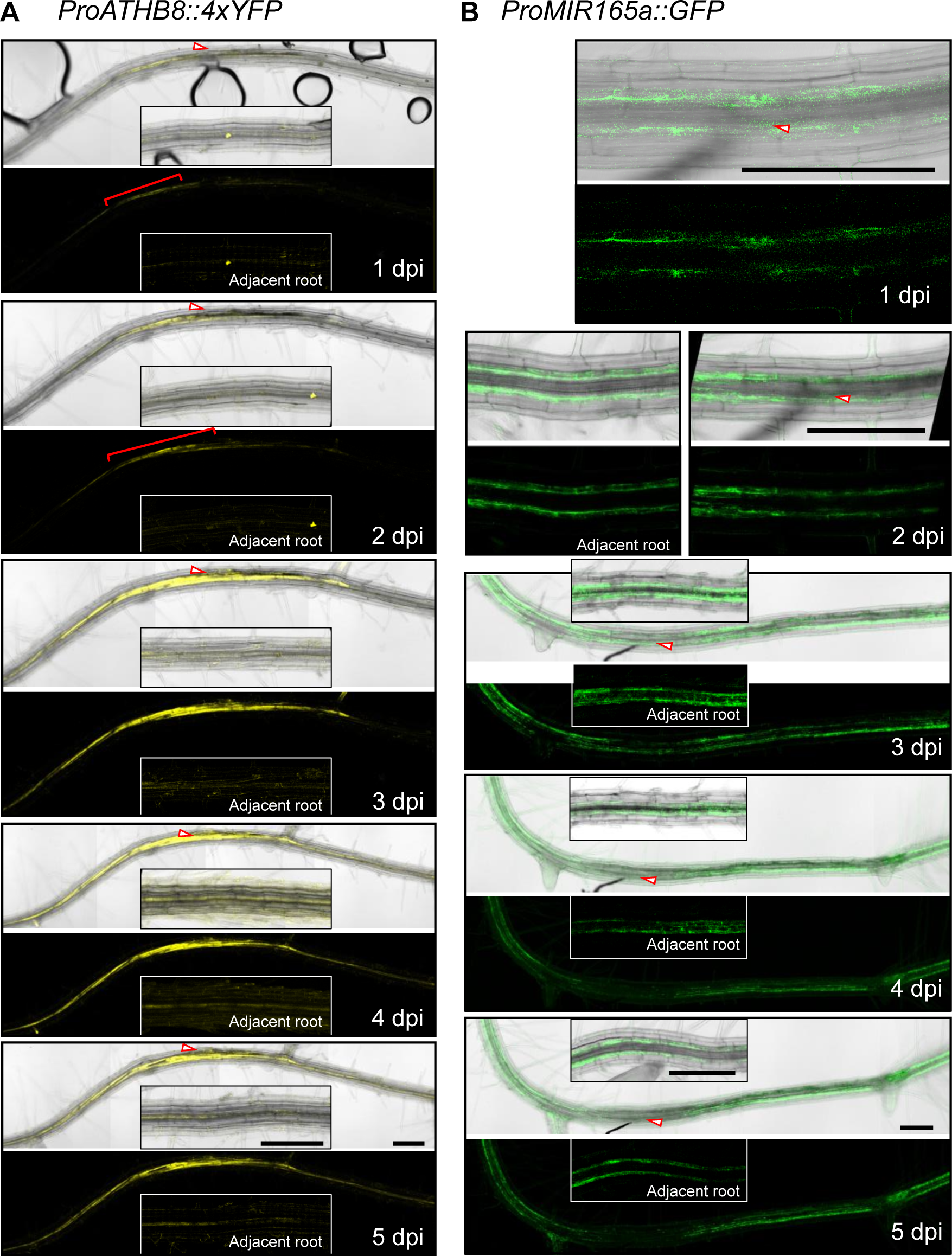
Expression of *ProATHB8::4xYFP* and *ProMIR165a::GFP* at BCN infection sites in *Arabidopsis* roots. **A.** Continuous monitoring of *ProATHB8::4xYFP* expression in early stages of BCN infection. Red brackets indicate increased YFP signal at early stages of BCN infection. Insets represent YFP signal of root segment about 850 μm away from the infection site. **B.** Continuous monitoring of *ProMIR165a::GFP* in early stages of BCN-infected roots. Left panel of 2 dpi, and insets of 3, 4, and 5 dpi panels represent GFP signal of root segment adjacent to the infection site. Red arrowhead, position of nematode. bar = 200 um.

To evaluate if *ATHB8* expression is regulated by CLE receptors, we introduced the *ProATHB8::GUS* transgene into the *clv* triple mutant by crossing. In the *clv* triple mutant, the transgene showed a similar expression pattern and expression level to that of the wild-type (Supplemental Figure S9A, B). Upon BCN infection, *ProATHB8::GUS* expression was also up regulated in the infection site of the *clv* triple mutant root, and the increasing levels were comparable to that of the wild-type (Figure 3F-J). The increased expression of *ProATHB8::GUS* in the mutant could be mediated by other CLE receptors, like BAMs and TDR (Guo et al., 2011; Guo et al., 2017; Crook et al., 2020). In addition, *ATHB8* is directly regulated by auxin response factor ARF5/MP (Donner et al., 2009). High expression of *ARF5/MP* after BCN infection could also induce the up-regulation of *ProATHB8::GUS* in the *clv* triple mutant in the infection site (Supplemental Figure S6) (Hewezi et al., 2014). In addition, given that the *ProATHB8::GUS* is unevenly expressed along the vasculature of root (Supplemental Figure S9A, B), and that *ProATHB8::GUS* expression varies greatly at the infection site, it is challenging to compare *ATHB8* expression level differences across genotypes. Our RNA sequencing and qPCR data consistently showed that *ATHB8* gene expression in the *clv* triple mutant infection site is only about 25% lower compared to that of wild-type (Figure 2C-D), and therefore, the GUS staining technique may not be sensitive enough to distinguish such a small difference in gene expression (Figure 3A-J).

### *MIR165a* Expression is Down-Regulated at the BCN Infection Site

*ATHB8*, as a member of the HD-ZIP III family of genes, is post-transcriptionally regulated by the *MIR165/166* family of microRNAs. *Arabidopsis* encodes two *MIR165* genes and seven *MIR166* genes, of which *MIR165a*, *MIR166a*, and *MIR166b* are expressed in roots, and only *MIR165a* and *MIR166b* promoters drive detectable GFP expression (Jung and Park, 2007; Carlsbecker et al., 2010). A previous study reported no significant difference of MIR165/166 expression in the BCN infection site in *Arabidopsis* roots (Hewezi et al., 2008a). To examine the response of *MIR165*/*6* to BCN infection, two transgenic lines, *ProMIR165a::GFP* and *ProMIR166b::GFP*, were used to track their expression in the infection site. In uninfected roots, *ProMIR165a::GFP* is expressed in the endodermal cells in both the root tip and mature root (Supplemental Figure S11A) (Carlsbecker et al., 2010). Upon BCN infection, expression of *ProMIR165a::GFP* signal is gradually reduced at the infection site (Figure 4B); whereas for *MIR166b*, GFP signal was only detected in the endodermal cells of root tips in both uninfected and infected plants (Supplemental Figure S11B). No signal was detected in the infection site up to five days post infection (Supplemental Figure S11C, D). Furthermore, in the RNAseq dataset, *AGO1* (*AT1G48410*), a gene encoding an ARGONAUTE protein that recruits MIR165/6 for target mRNA cleavage (Baumberger and Baulcombe, 2005), is downregulated; while *AGO10* (AT5G43810), which encodes an ARGONAUTE that specifically sequesters *MIR165/6* for degradation (Zhu et al., 2011; Yu et al., 2017, 2021), is up-regulated (Supplemental Figure S12). These results point to a reduced *MIR165/6* level and activity at the BCN infection site.

To test if the MIR165/6 activity is indeed reduced, the *U2::MIR165/6-GFP MIR165/6 sensor* line, which harbors a MIR165/6 target sequence at the 3’UTR of ER-GFP construct (Carlsbecker et al., 2010), was used to monitor the MIR165/6 activity at the early stage of BCN infection. In this system, high *MIR165/6* activity is represented by low GFP expression (Carlsbecker et al., 2010). In uninfected roots, high GFP expression was observed in the vasculature, representing low levels of *MIR165/6* activity in corresponding tissue (Supplemental Figure S13A). After BCN infection, we did not observe a clear increase in GFP signal, due to the preexisting high level of GFP expression (low MIR165/6) (Supplemental Figure S13B); however, the GFP expression domain was slightly expanded at the infection site (Supplemental Figure S13B), probably due to reduced MIR165 expression in the endodermis cells (Figure 4B).

### Loss of Function of *ATHB8* and *ATHB15*/*CNA* Does Not Affect BCN Infection

To test if *ATHB8* plays any role in syncytium formation, we obtained single and double knockout mutants of *ATHB8* and its closest homology *ATHB15*/*CNA*, *athb8-11* and *cna-2* respectively (Prigge et al., 2005; Carlsbecker et al., 2010). In our growth conditions, young seedlings of *athb8-11*, *cna-2*, and *athb8-11 cna-2* mutants all showed similar root and shoot growth and development compared to that of the wild-type control Col *er-2* (Figure 5A-C). Comparable numbers of BCN juveniles were able to penetrate into the roots of these mutants and wild-type plants (Figure 5D). The number of adult females that developed on these mutants’ roots were also comparable to that of wild-type plants (Figure 5E), and no significant differences were observed in syncytium size in any of these mutants compared to those developed on wild-type plants (Figure 5F). These results indicated that knocking out *ATHB8* and *ATHB15*/*CNA* genes does not affect syncytium formation and syncytium expansion in the *Arabidopsis* root, probably due to high redundancy of HD-ZIP III family genes in mediating vascular cell differentiation, and that all five members were up-regulated at the BCN infection site (Prigge et al., 2005; Carlsbecker et al., 2010)(Supplemental Figure S8).

**Figure 5.**
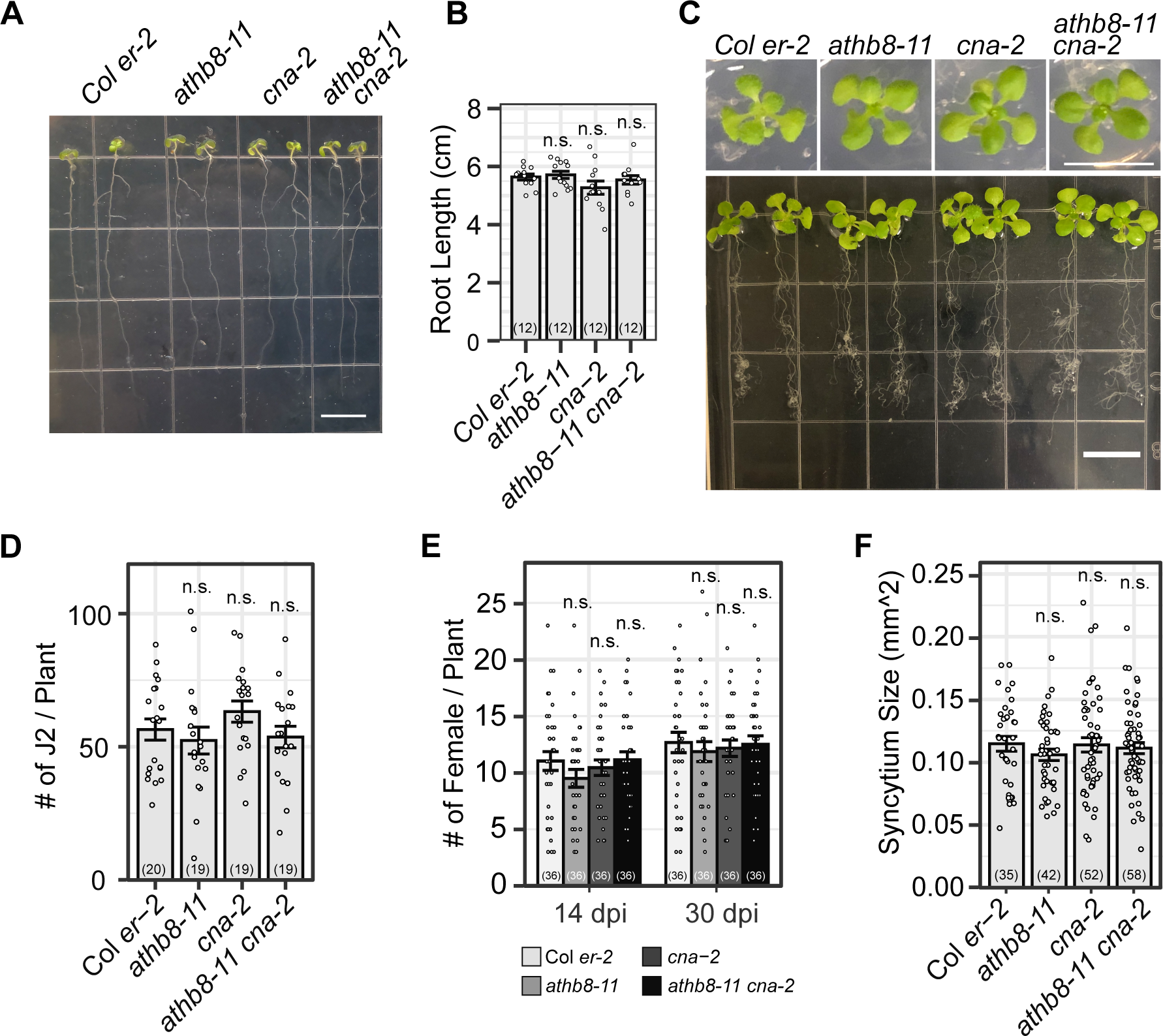
Loss of function of *ATHB8* and its closest homologue *ATHB15/CNA* do not affect BCN infection or syncytia development in *Arabidopsis*. **A.** 7-day-old seedlings of *athb8-11*, *cna-2*, and *athb8-11 can-2* grown on a vertical plate. bar = 1 cm **B.** Measurement of root length shown in (**A**). **C.** 14-day-old seedlings grown in 12-well plates. Top panel, top view of representative seedlings. Bottom panel, root system of representative 14-day-old seedlings grown in 12-well plates and scooped for photograph. bar = 1 cm. **D.** BCN penetration rate on the tested mutants, counted at 3-5 days post inoculation. **E.** Number of adult females developed on the tested mutants. **F.** Syncytium size developed on the tested mutants. All bar graphs represent mean ± SE. Dots represent each individual measurement. Samples sizes were shown in parentheses in each bar. n.s., not significant compared to control group with Student’s t-test. Data shown in **D-F** were repeated twice with similar results.

### Knock Down of the HD-ZIP III Gene Family Reduced BCN Infection Rate & Syncytium Development

To overcome functional redundancy among HD-ZIP III genes, higher order mutants would be needed for BCN infection assays. However, vascular development defects in these high order mutants, especially quadruple and quintuple HD-ZIP III mutants results in severe root growth phenotypes, making them unsuitable for assessing BCN infection phenotypes (Prigge et al., 2005; Carlsbecker et al., 2010; Smetana et al., 2019). To further evaluate the role of HD-ZIP III genes in cyst nematode infection, we adapted the estradiol-based inducible gene expression system to the cyst nematode infection assay. Two *MIR165a*-inducible expression lines, *Pro35S::XVE>>MIR165a* and *ProCRE1::XVE>>MIR165a* (hereafter designated as *Pro35SiMIR165a* and *ProCRE1iMIR165a*) (Muller et al., 2016; Smetana et al., 2019), were used to knock down HD-ZIP III gene expression at the time of nematode infection. Upon 48 hours of induction by 5 μM 17-beta-estradiol, expression of all members of the HD-ZIP III gene family in the *Pro35SiMIR165a* line were significantly suppressed (Figure 6A). Suppression of gene expression in *ProCRE1iMIR165a* were less severe compared to that of the *Pro35SiMIR165a* line, yet expression of all members of HD-ZIP IIIs except for *REV* were significantly decreased (Figure 6A). Increasing estradiol concentration to 10 μM, or prolonging the incubation time to 72 hours, did not further reduce target gene expression (Figure 6A). In fact, when estradiol induction was extended beyond three days, the expression levels of all HD-ZIP III genes gradually recovered, highlighting the transient effect of estradiol induction (Figure 6B). This expression profile, however, is advantageous for nematode infection assays, as the disruption of the target gene can be timed with nematode infection, and any adverse effects of target gene knock-down on plant development can be largely minimized (Figure 6B). Based on this information, 5 μM estradiol was used for *MIR165a* induction on 12-day-old seedlings and BCN was inoculated 48 hours after induction. Estradiol treatment did not affect BCN activity, as estradiol and DMSO treated *Pro35SiGUS* lines showed similar penetration and infection rate, as well as female associated syncytium size (Figure 6C-E). These numbers were also comparable to nematode infection on wild-type plants without any treatment (Figure 5D-E). In either *Pro35SiMIR165a* or *ProCRE1iMIR165a* line, nematode penetration rates were not significantly affected by estradiol treatment, counted at five days post inoculation (Figure 6C). Whereas the development of adult females on both inducible lines was strongly suppressed (Figure 6D), indicating that HD-ZIP III TFs do play essential roles in syncytium formation during BCN infection. Interestingly, although expression levels of HD-ZIP III genes were only slightly decreased in the *ProCRE1iMIR165a* line, and target gene expression recovered much faster compared to that of the *Pro35SiMIR165a* line (Figure 6A, B), a comparable level of suppression of female development to that of the *Pro35SiMIR165a* line was observed (Figure 6D). In addition, knocking down HD-ZIP IIIs by inducible expression of *ProCRE1iMIR165a* significantly reduced average syncytia size (Figure 6E), whereas in the *Pro35SiMIR165a* line, although the average size of syncytia was consistently smaller, the change was not statistically significant compared to the control (Figure 6E). This difference in syncytia size may be attributed to the different root cell-type expression patterns and nematode regulation of the 35S (constitutive) and *CRE1* (stele) promoters, and suggested that suppression of HD-ZIP III expression in the stele is sufficient to suppress syncytium formation and expansion by BCN.

**Figure 6.**
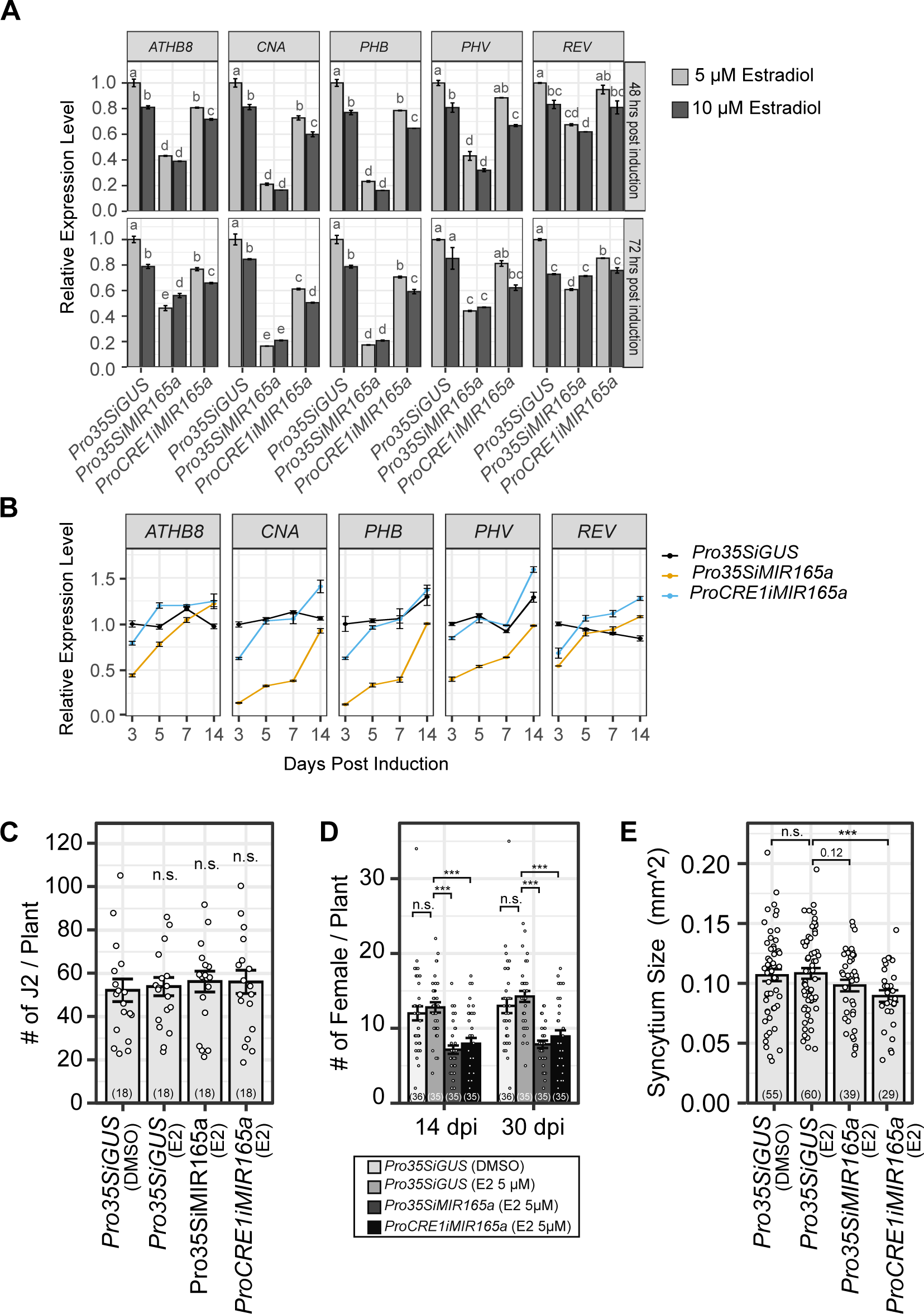
Inducible expression of *MIR165a* suppresses BCN infection in *Arabidopsis*. **A.** Induction of *MIR165a* suppressed expression of HD-ZIP III TFs in 12-well plates. Top panel, 48 hours post-induction. Bottom panel, 72 hours post-induction. Letter above each bar represents statistical group of Turkey’s HSD test following ANOVA analysis. **B.** Expression of HD-ZIP III TFs 3-14 days post induction of *MIR165a* with 5 μM estradiol (E2). Effect of *MIR165a* induction attenuates over time. **C.** Suppression of HD-ZIP III TFs did not affect nematode penetration into the root. Penetration rate counted at 5 dpi. **D.** Suppression of HD-ZIP III TFs suppressed BCN female development on *Arabidopsis* roots. **E.** Suppression of HD-ZIP III TFs resulted in smaller syncytia size. Bar graph represents mean ± SE. Dots represent each individual measurement. Sample sizes were shown in parentheses in each bar. Statistical tests were performed with student’s t-test, n.s., not significant; **, p < 0.01; ***, p < 0.001. Data shown in (**C-D)** were repeated four times with similar results. Data shown in (**E)** was repeated twice with similar results. E2, estradiol.

### Overexpression of *ATHB8* Does Not Induce Hypersusceptibility to BCN

Next, we tested if overexpression of *ATHB8* would result in cyst nematode hyper-susceptibility. *ATHB8* overexpression lines were created by crossing *ProANT::XVE>>ATHB8d-YFP* (*ProANTiATHB8d-YFP*), which expresses *MIR165* resistant *ATHB8* (*ATHB8d*) using the *ANT* promoter (Smetana et al., 2019), with *Pro35S::XVE>>GUS* (*Pro35SiGUS*) and *ProG1090::XVE>>GUS* (*ProG1090iGUS*) lines. F1 of these crosses, designated as *Pro35SiATHB8d-YFP* and *ProG1090iATHB8d-YFP* respectively, contain constructs from both parents thus the *ATHB8d-YFP* transgene can be activated by both the *ANT* promoter and *35S*/*G1090* promoters, resulting in ubiquitous overexpression. Indeed, in the root of F1 plants, *ATHB8d-YFP* expression domains were much expanded compared to that of the parent line *ProANTiATHB8d-YFP* (Figure 7A). qPCR showed that in all lines, expression level of *ATHB8* was highly induced three days post induction, up to 70 times higher in the *ProG1090iATHB8-YFP* line, and 10 times higher in *Pro35SiATHB8d-YFP* and *ProANTiATHB8d-YFP* lines (Figure 7B). Consistent with previous observations (Figure 6B), the levels of induced gene expression declined over time (Figure 7B). Expression of other HD-ZIP III TF family members was not substantially affected by *ATHB8d* overexpression (Figure 7B). In all tested lines, overexpression of *ATHB8d* did not affect penetration of nematodes into the root (Figure 7C) or average syncytia size induced by nematodes (Figure 7E). In *35S* and *ANT* promoter driven lines, which have relatively low overexpression levels, female development was not significantly affected (Figure 7D). However, in the *ProG1090iATHB8d-YFP* line, nematode female development was strongly suppressed (Figure 7D). This distinct phenotypic difference in female development between *G1090* driven and *35S*/*ANT* driven lines may be due to the extremely high *ATHB8d-YFP* expression level and/or expanded expression domain of the *G1090* promoter compared to the other two lines (Figure 7A, B). Induction of *ProG1090iATHB8d-YFP* in younger (5-day old) seedlings caused severe developmental defects in both roots and shoots when grown on vertical plates, regardless of BCN infection (Supplemental Figure S14), while *ProANTiATHB8d-YFP* and *Pro35SiATHB8d-YFP* did not show a clear phenotype (Supplemental Figure S14). However, developmental defects of the *ProG1090iATHB8d-YFP* line were not obvious for seedlings used in 12-well plate infection assays (Supplemental Figure S15), where seedlings were much older (12-day) and had more developed root systems that masked the subtle root growth phenotypes observed on vertical plates (Supplemental Figure S14). Nevertheless, the defects in root development were unlikely to directly contribute to reduced female development, as the root system has not been significantly affected by *ATHB8d-YFP* overexpression at the point of BCN inoculation, and comparable numbers of nematodes were able to penetrate into the root system in all lines (Figure 8C). These results suggested that overexpression of *ATHB8* is not sufficient to induce BCN hyper-susceptibility. Instead, the balance of stem cell organizer cell specification and its further differentiation might be important for syncytial cell transition.

**Figure 7.**
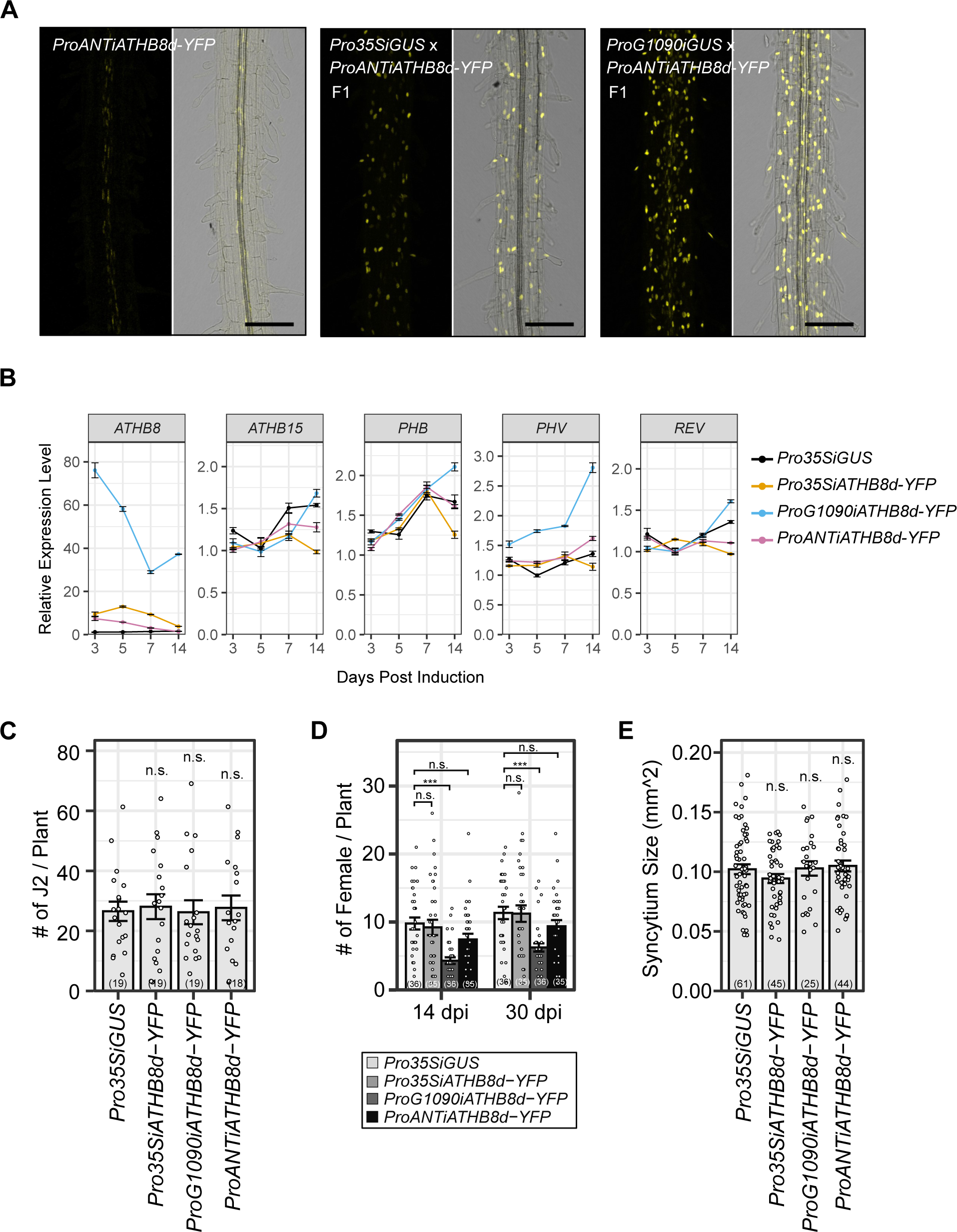
Effect of overexpressing *ATHB8* on cyst nematode infection in *Arabidopsis*. **A.** Expanded *ATHB8d-YFP* expression domain after introduction to *Pro35SiGUS* and *pG1090iGUS* lines. 5-day-old seedlings were induced with 5 μM estradiol for 24 hours before imaging. bar = 100 μm. **B.** Expression of HD-ZIP III genes in 12-well plates after estradiol (5 μM) induction. **C-E.** Effect of ATHB8d overexpression on BCN penetration (**C**), female development (**D**), and syncytia size (**E**) in *Arabidopsis* root. Bar graph represents mean ± SE. Dots represent each individual measurement. Sample sizes were shown in parentheses in each bar. Statistical tests were performed with student’s t-test, n.s., not significant; ***, p < 0.001. Data shown in **C-E** were repeated twice with similar results. Data shown in **(E)** was repeated twice with similar results.

**Figure 8.**
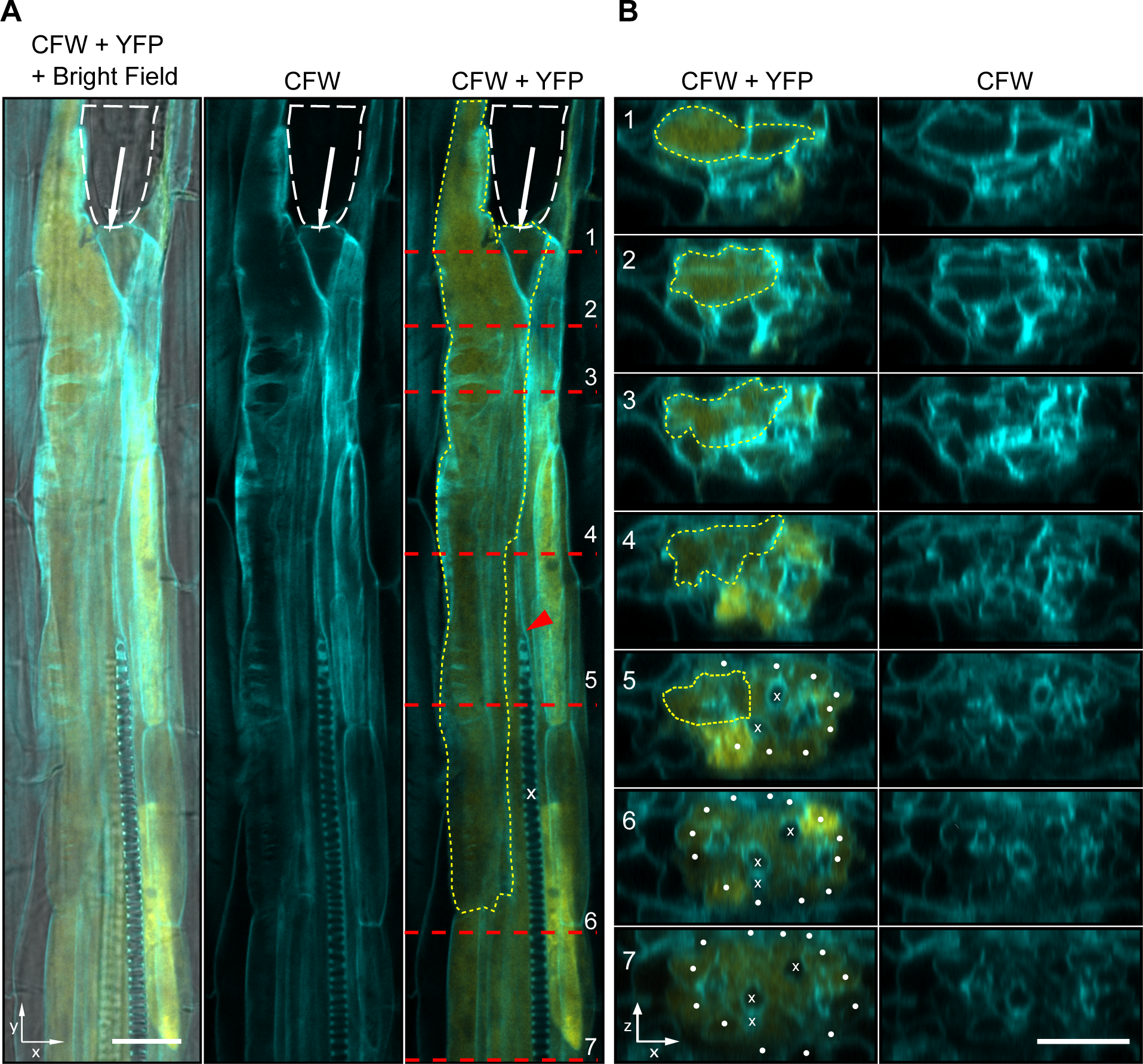
Expression of *ProATHB8::4xYFP* in the BCN induced syncytium at 1 dpi. **A.** A confocal optical section of developing syncytium with *ProATHB8::4xYFP* expression. **B.** Optical cross section of positions shown in (**A**). White dashed line, outline of nematode head. White arrow, position of the stylet. Red dashed line, positions of cross section shown in (**B**). Yellow dashed line, outline of the syncytium. x, xylem cells. Red arrowhead, disrupted xylem differentiation. White dot, pericycle cells. CFW, calcofluor white staining of cell wall. YFP, yellow fluorescent signal.

### Auxin Level and *ATHB8* Expression are Elevated in Neighboring Cells of the Developing Syncytium

Considering the finding that *ATHB8* expression is upregulated at the periphery of syncytium (Figure 3; Supplemental Figure S10B), coupled with prior studies demonstrating *ATHB8* can be directly regulated by auxin through ARF5/MP, an auxin responsive factor essential for vascular development (Donner et al., 2009), and that auxin was proposed to function in conditioning syncytium periphery cells for syncytial integration (Grunewald et al., 2009; Lee et al., 2011), we reasoned that *ATHB8* may function downstream of auxin in conditioning cells for incorporation into the syncytium. To explore this hypothesis, we carried out a detailed analysis of the spatial distribution of auxin response and *ATHB8* promoter activities at early stages of syncytium formation. Indeed, at 24 hpi (hours post inoculation), both *ProATHB8::4xYFP* and *DR5::4xYFP* were seen in the developing syncytia (Figure 8A, B; Supplemental Figure S16A - B). By this stage, the syncytia were already well developed with extensive cell wall modifications and fused cytoplasm (Figure 8A - B, Supplemental Figure S16A - B), and higher YFP signal was frequently observed in cells neighboring syncytial cells (Figure 8A - B; Supplemental Figure S16A - B). Nevertheless, to determine if auxin level or *ATHB8* expression peaked earlier than this is technically challenging, as it is difficult to determine if the nematode has begun feeding yet, or pinpoint the initial syncytial cell without some visible cell wall modification or a reliable syncytium specific marker. In both *ProATHB8::4xYFP* and *DR5::4xYFP* lines, the YFP signal was also observed in pericycle cells incorporated into the syncytium (Supplemental Figure S16B), or pericycle cells near the syncytium (Figure 8B). These two transgenes are typically not expressed in pericycle cells of uninfected roots, except where lateral roots initiate (Torres-Martinez et al., 2022). At 3 dpi, YFP signal in the syncytium was barely detectable in the syncytial cell itself, but observed primarily in cells neighboring the developing syncytium (Figure 9A-C; Supplemental Figure S17A - C). These results showed that indeed, auxin and *ATHB8* levels first accumulate in the early-stage syncytial cells and then expression shifts to adjacent cells of the developing syncytium (Grunewald et al., 2009), implying that auxin signaling and *ATHB8* may function in promoting the transition of a host cell into a syncytial cell.

**Figure 9.**
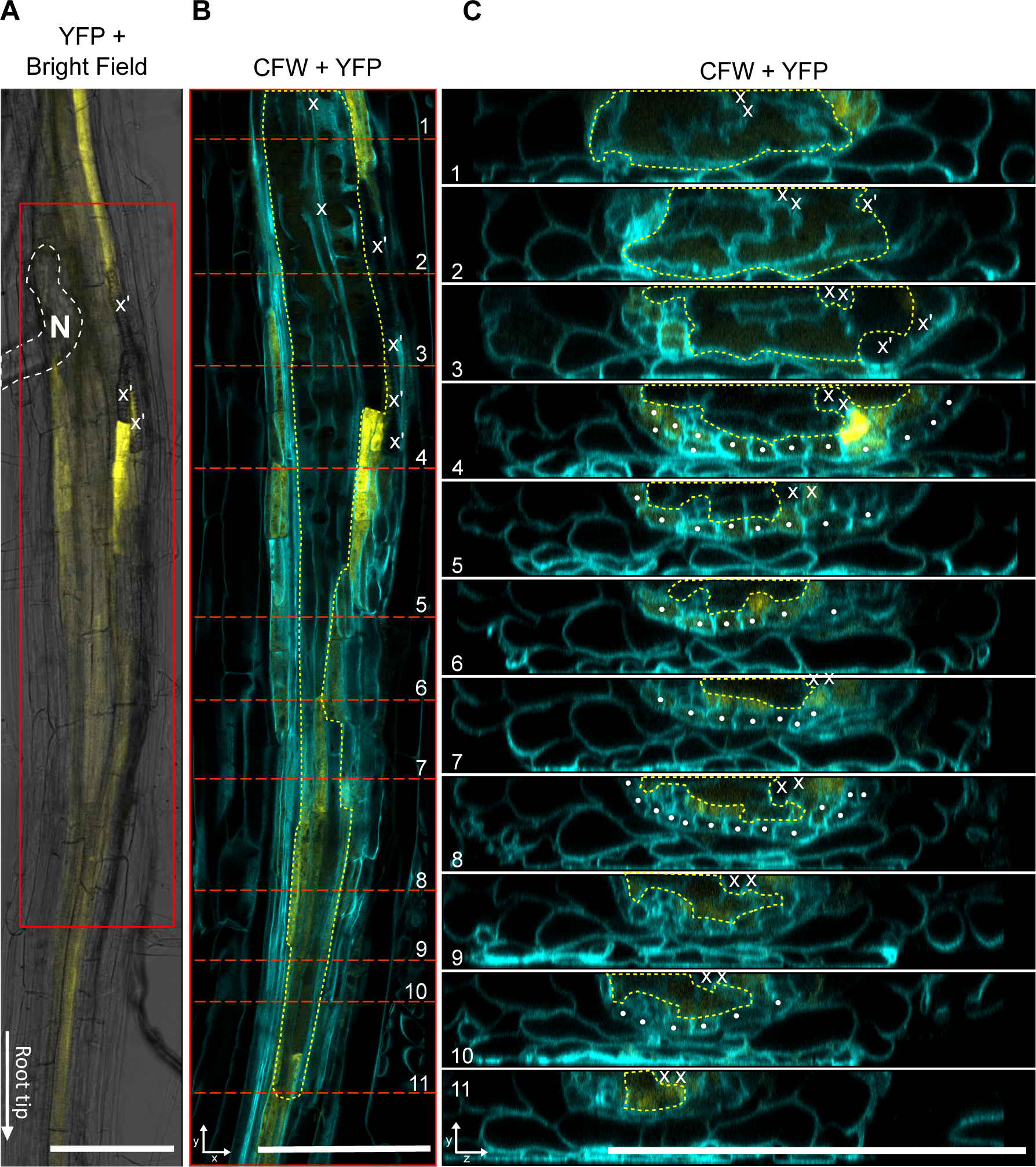
Distribution of *ProATHB8::4xYFP* activity at BCN infection site at 3 dpi. White dashed line, outline of nematode. Red box, zoomed in portion in (**B**). Red dashed lines in (**B**), position of optical cross sections shown in (**C**). Yellow dashed line, outline of the syncytium. x, xylem cells. x’, ectopic xylem cells. White dot, pericycle cells. CFW, calcofluor white staining of cell wall. YFP, yellow fluorescent signal.

## Discussion

### Gene Transcription Profiles of CLE Receptor Mutant and Wild-type Roots Converge Upon Cyst Nematode Infection

Cyst nematode CLE-like peptide effectors were the first stylet-secreted effectors found to mimic host peptide hormones to modulate plant developmental programs for parasitic success (Wang et al., 2005; Lu et al., 2009; Wang et al., 2011). Impairing CLE receptors in host plants attenuates the CLE signaling pathway and increases host resistance to cyst nematode infection (Replogle et al., 2011; Replogle et al., 2012; Chen et al., 2015; Guo et al., 2015; Guo et al., 2017).

*Arabidopsis* CLE receptors CLV1, CLV2, and RPK2, and their orthologs in soybean and potato, play an important role in mediating CN infection (Replogle et al., 2012; Guo et al., 2017; (Chen et al., 2015). Simultaneously knocking out or knocking down the three receptors results in lower nematode infection rate and smaller syncytia size (Replogle et al., 2012; Guo et al., 2017; (Chen et al., 2015). Like in other classical CLE-RLK-WUS/WOX signaling modules, the *WOX4* gene, which functions in maintaining vascular cambial activity (Hirakawa et al., 2010; Ji et al., 2010; Suer et al., 2011; Etchells et al., 2013), is up-regulated by nematode infection through A- and B-type CLE receptors (Guo et al., 2017). Nevertheless, knocking out *WOX4* does not yield similar levels of BCN resistance as the *clv1-101 clv2-101 rpk2-5* triple CLE receptor mutant, indicating other CLE downstream factors are at play during BCN infection (Guo et al., 2017). Nevertheless, trying to isolate such downstream factors by directly comparing BCN-infected wild-type and triple CLE receptor mutant transcriptomes proved difficult, as most DEGs responding to BCN infection in the wild-type were also differentially expressed in the mutant (Figure 1C; Supplemental Figure S3A). Moreover, BCN infection further reduced the difference in the gene transcription profiles between the mutant and wild-type roots compared to uninfected roots (Figure 2E; Supplemental Figure S2B, C; S3C; S4A, B), whereas the HsCLE2p treatment widened the difference of gene expression profiles between the two (Figure 2F; Supplemental Figure S2B, C; S3D; S4C, D). These results suggested that although loss of CLE receptors suppressed nematode infection, once the initial barrier was overcome and syncytium development was initiated, the nematodes were capable of inducing an attenuated, yet similar gene expression profile sufficient for syncytium establishment and maintenance, albeit with some size restriction that may compromise feeding and lead to developmental delays (Replogle et al., 2011; Replogle et al., 2012). This convergence of gene expression profiles of mutant and WT roots responding to BCN infection also highlights the robustness of nematode effectors in co-opting plant developmental programs. The indifference of syncytia transcriptomes between the triple CLE receptor mutant and wild-type could be due to secretion of a mixture of A- and B-type CLE peptide mimics (Wang et al., 2011; Guo et al., 2017), that could signal through multiple CLE receptors and function redundantly or in parallel to bypass these impaired CLE receptors (Fletcher, 2020). Moreover, modulation of other signaling pathways, like auxin and cytokinin, by other nematode effectors, could also compensate for the lack of these CLE receptors, given that these signaling pathways are highly intertwined with CLE signaling in regulating vascular cambium activity (Cammarata et al., 2019; Willoughby and Nimchuk, 2021; (Fischer et al., 2019). For example, *WOX4*, the infamous gene in regulating vascular cambium activity downstream of CLE signaling, is also responsive to auxin and cytokinin, both of which function in modulating a complex transcriptional network to regulate vascular cambium activity (Suer et al., 2011; Brackmann et al., 2018; Zhang et al., 2019; Smit et al., 2020; Ye et al., 2021). During cyst nematode infection, an overdrive of these hormone signaling pathways could contribute to the up-regulation of CLE downstream genes like *WOX4*, thereby masking effects of attenuated CLE signaling on their expression.

For this reason, we primarily used the HsCLE2 peptide effector treated samples to identify potential genes downstream of nematode CLE signaling pathways. 147 and 176 genes were identified to be positively or negatively regulated, respectively, by CLV1, CLV2, and RPK2 CLE receptor-like-kinases in response to BCN infection (Figure 2A, B; Supplemental Figure S7A, B; Supplemental Dataset S4-5). These genes are likely to be directly regulated by CLE peptide signaling, rather than being a secondary effect of HsCLE2-induced changes in vascular development, as they responded relatively quickly (3 hours) to a relatively low dose (5 μM) of HsCLE2p treatment, on par with the response of the *WOX4* gene (Supplemental Figure S1A). The list of genes positively regulated by CLE signaling was particularly interesting to us. Not only because *WOX4*, the only known marker gene for CLE signaling in vascular development and cyst nematode infection, was included in the list, but because this list also included a number of genes known to be involved in vascular development or cell fate determination, such as *BAM2*, *SERK1*, *PXC1/2*, *ANT*, *CYCD3;1*, *LBD4*, *OPS*, and *ATHB8* (Supplemental Dataset S4) (Baima et al., 2001; Truernit et al., 2012; Wang et al., 2013; Collins et al., 2015; Randall et al., 2015; Zhang et al., 2016; Breda et al., 2019; Smetana et al., 2019; Smit et al., 2020; Fan et al., 2021). A few of these genes were also reported to be regulated by CLE signaling. For example, *ATHB8* had been shown to be up-regulated by CLE peptide treatment (Whitford et al., 2008; Hirakawa et al., 2010); whereas *LBD4* was identified as a part of the *TMO6*-*WOX14*-*LBD4* feedforward network regulated by CLE and auxin signaling in regulating vascular development (Smit et al., 2020). In addition to being potentially regulated by CLE signaling, a few of these genes were also involved in syncytium formation through other hormone signaling pathways, like the *PIN3* and *PIN4* genes that were proposed to distribute auxin to syncytium lateral cells for its expansion (Grunewald et al., 2009), and the *CKX6* and *CKX7* genes that were shown to be up-regulated at BCN infection sites and may modulate infection through cytokinin catabolism (Dowd et al., 2017). As potential targets of CLE signaling pathways, these genes may represent intersections between CLE and other phytohormone signaling during syncytium formation (Gheysen and Mitchum, 2019).

### *ATHB8* Gene Expression is Regulated by CLE Peptide Signaling and BCN Infection

*ATHB8* belongs to the HD-ZIP III family of transcription factors, which play essential roles in vascular development (Ramachandran et al., 2017). *ATHB8* has been shown to be up-regulated by 10 μM CLE41p/TDIF (B-type) within one day of treatment, and co-application of CLE6p (A-type) with CLE41p accelerated *ATHB8* up-regulation to 3 hours (Whitford et al., 2008). Here, we showed that application of 5 μM HsCLE2p, a peptide effector mimic of CLE6p (Wang et al., 2011), induced significant up-regulation of *ATHB8* within 3 hours of treatment (Figure 2C-D), a timeline on par with co-treatment of A- and B-type CLEs (Whitford et al., 2008). Like the *tdr-1* mutant, the *clv1-101 clv2-101 rpk2-5* mutant eliminated the response of *ATHB8* to corresponding CLE peptide treatments (Guo et al., 2017) (Figure 2C-D), suggesting corresponding receptors are required for A- or B-type CLE-induced *ATHB8* expression at this time point. One potential way HsCLE2p/CLE6p could promote *ATHB8* expression is by up-regulating *CLE41* gene expression (Supplemental Figure S2A). The CLE41p/TDIF-TDR module has been shown to signal through BIN2/BIL1 to phosphorylate members of the closely related auxin response factors (ARF), ARF5/MP, ARF7, and ARF19, to release them from the inhibitory IAA proteins (Cho et al., 2013; Han et al., 2018). These ARF factors could then activate *ATHB8* gene expression by directly binding to its promoter (Donner et al., 2009). Nevertheless, it is also likely that HsCLE2p/CLE6p has its own pathway to regulate *ATHB8* expression, given that co-application of HsCLE2p/CLE6p greatly enhanced CLE41p/TDIF-induced *ATHB8* expression (Whitford et al., 2008).

*ATHB8* expression was also strongly up-regulated by BCN infection (Figure 2C-D; Figure 3A-E, Figure 4A). Multiple mechanisms could explain the high accumulation of *ATHB8* transcripts at BCN infection sites. First, the *ATHB8* promoter was highly activated at BCN infection sites (Figure 3A-E; Figure 4A; Figure 8; Figure 9). Besides CLE signaling, elevated auxin level at the infection site could also directly contribute to the activation of *ATHB8* promoter through transcription factors like ARF5/MP (Supplemental Figure S16, S17) (Karczmarek et al., 2004; Grunewald et al., 2008; Donner et al., 2009; Abril-Urias et al., 2023), which itself was also highly up-regulated at the BCN infection site (Supplemental Figure S6) (Hewezi et al., 2014).

Interestingly, although *ATHB8* promoter activity was slightly elevated in early syncytium compared to adjacent root (Figure 8), its activation was much more prominent in cells neighboring the syncytium rather than in the syncytium itself (Figure 8). By the late J2 or early J3 stage of BCN development (3 – 5 dpi), *ATHB8* expression was barely detectable in the syncytium, but high in some of the neighboring cells (Figure 3A; Figure 9A, B). Previously, it was shown that CN CLE effectors are re-secreted into the apoplast after being injected into the host cell cytoplasm, and therefore could potentially activate CLE signaling pathways of adjacent cells (Wang et al., 2010a; Wang et al., 2010b; Wang et al., 2021). In addition, auxin was also proposed to be first concentrated in the initial syncytium cell, and then distributed laterally to adjacent cells to facilitate syncytium expansion (Grunewald et al., 2009; Lee et al., 2011). Both of these factors may contribute to the lateral shift of the *ATHB8* promoter activation domain.

Second, lowered *MIR165/166* level also contributes to high *ATHB8* accumulation at the BCN infection site. Of the two *MIR165/166* promoters that drove detectable GFP signal in *Arabidopsis* roots (Carlsbecker et al., 2010; Vaten et al., 2011), *ProMIR165a::GFP* was down-regulated at the infection site (Figure 4B), and *ProMIR166b::GFP* was not detectable at the infection site or corresponding uninfected root segments (Supplemental Figure S11B-D), pointing to a reduced *MIR165/166* expression level at the infection site. *MIR165/166* genes expression requires transcription factor SHOOT-ROOT (SHR) to move from stele cells to the endodermis layer, through plasmodesmata, and its subsequent nuclear localization (Helariutta et al., 2000); (Nakajima et al., 2001; Gallagher et al., 2004; Cui et al., 2007; Carlsbecker et al., 2010; Vaten et al., 2011; Long et al., 2015). Although *SHR* gene expression was not significantly affected during BCN infection (Supplemental Figure S12), the movement of SHR from stele cells to the endodermis may be restricted, considering that the syncytium is largely symplastically isolated from surrounding root cells, and very few plasmodesmata were found in the outer wall of syncytium (Böckenhoff et al., 1996; Grundler et al., 1998; Hofmann et al., 2007; Hofmann et al., 2010); (Bockenhoff and Grundler, 1994) except for plasmodesmata connections with neighboring phloem cells (Hoth et al., 2005; Hofmann and Grundler, 2006; Hoth et al., 2008).

Symplastic isolation could also limit the movement of *MIR165/166* from the endodermis to the syncytial cell and reduce its level in the syncytium (Vaten et al., 2011). Furthermore, the *AGO1* gene, which encodes an ARGONAUTE that recruits microRNAs for mRNA cleavage (Baumberger and Baulcombe, 2005), was down-regulated (Supplemental Figure S12); while the gene encoding AGO10, which specially sequesters *MIR165*/*166* and promotes their degradation (Zhu et al., 2011; Yu et al., 2017, 2021), was up-regulated (Supplemental Figure S12). Collectively, these mechanisms could regulate *ATHB8* expression level in and around syncytium and result in high *ATHB8* transcript accumulation (Figure 2C, D).

Co-existence of multiple mechanisms in regulating *ATHB8* expression means that disrupting CLE signaling alone would not be sufficient to completely abolish BCN-induced *ATHB8* up-regulation, like in the situation of HsCLE2p treatment (Figure 2C-D). Thus, it is not surprising that *ATHB8* was also highly up-regulated at the BCN infection site in the *clv1-101 clv2-101 rpk2-5* triple mutant (Figure 2C-D; Figure 3F-G). Nevertheless, the expression level of *ATHB8* in the mutant did not reach to the same level as the wild-type infection sites (Figure 2C-D), suggesting that CLE signaling is indeed required for optimal *ATHB8* expression during syncytium formation.

### HD-ZIP III Transcription Factors Function in Syncytium Formation

*ATHB8*, along with other HD-ZIP III family members, promote xylem differentiation during vascular development in a dose-dependent manner (Baima et al., 2001; Prigge et al., 2005; Carlsbecker et al., 2010). While single or double mutants of HD-ZIP III family members have no or very minor defects in xylem differentiation, higher order mutants exhibit progressively more severe phenotypes as more members are knocked out (Carlsbecker et al., 2010). Ultimately, the *athb8-11 cna-2 phb-13 phv-11 rev-6* quintuple mutant results in the complete loss of xylem axis cells (Carlsbecker et al., 2010), indicating all five genes are involved in xylem differentiation.

Functional redundancy among HD-ZIP III members may also be present in syncytium formation. All five members of HD-ZIP III genes were up-regulated by BCN infection (Supplemental Figure S8), and knocking out *ATHB8* and its closest homologous *ATHB15*/*CNA* was not sufficient to reduce the rate of BCN infection or average syncytium size (Figure 5E-F), indicating other HD-ZIP III members could compensate for the loss of function of these two genes in syncytium formation. Consequently, knocking down HD-ZIP III genes by inducible ectopic expression of *MIR165a* dramatically reduced the rate of successful BCN infection without affecting nematode penetration (Figure 6C-D). It is worth noting that expression of all five HD-ZIP III genes were still detectable after estradiol induction (Figure 6A), and that their expression levels gradually recovered over time after BCN inoculation (Figure 6B). Thus, if it were possible to knock out the activity of these genes locally in response to infection, syncytium establishment may be completely blocked, given the indispensable role of HD-ZIP III genes in vascular patterning (Carlsbecker et al., 2010). Interestingly, despite the fact that the *ProCRE1iMIR165a* line showed a relatively weaker suppression, and much faster recovery, of gene expression of all five HD-ZIP III members compared to the *35SiMIR165a* line (Figure 6A-B), it had a similar level of BCN suppression compared to that of the 35S driven line (Figure 6D), and an even greater effect on syncytium size (Figure 6E). These results may be due to the fact that the *CRE1* promoter is expressed at the infection site up to 5 dpi while 35S promoter is down-regulated as early as 3 dpi (Goddijn et al., 1993; Siddique et al., 2015). Alternatively, it could also mean that moderate suppression of HD-ZIP III gene expression in the stele, even within a short period of time at the beginning of BCN infection, is sufficient to suppress syncytium formation, as more nematodes failed to establish or were severely delayed in development, highlighting the importance of HD-ZIP III transcription factors in the initial establishment of the syncytium. The effect of HD-ZIP III gene knock-down on syncytium size was not as strong as its effect on initial syncytium establishment (Figure 6D-E), nevertheless, their involvement in syncytium expansion is still likely. In *Arabidopsis*, the BCN-induced syncytium reaches its maximum size at about 10 days after infection (Golinowski et al., 1996; Urwin et al., 1997). Given the fast recovery of HD-ZIP III gene expression, measuring syncytium size at 15 days post inoculation would give nematodes that overcame the initial difficulty in syncytium formation, and nematodes that started feeding late, time to catch up on syncytium expansion. Even so, both *ProCRE1iMIR165a* and *35SiMIR165a* showed consistent reduction in syncytium size (although it’s not statistically significant for the *35SiMIR165a* line) (Figure 6E). Overall, these results suggest that HD-ZIP III family transcription factors are important for initial syncytium establishment, and are also likely to be involved in syncytium expansion.

Recently, Smetana et al (2019) proposed that high auxin, and consequent expression of HD-ZIP III genes, promoted xylem identity and cellular quiescence in *Arabidopsis* root vasculature (Smetana et al., 2019). These xylem-identity cells serve as stem cell organizers, as they induce adjacent vascular cambial cells to divide and function as stem cells (Smetana et al., 2019).

Knocking down or knocking out HD-ZIP III genes forces xylem parenchyma cells and cells with xylem identity to re-enter mitosis, causing excessive cell proliferation (Carlsbecker et al., 2010; Smetana et al., 2019), while ectopic expression of ATHB8d in stem cells induced ectopic stem cell organizers and caused a vascular pattern shift (Smetana et al., 2019). Strong suppression of BCN infection in HD-ZIP III knock down lines suggests that HD-ZIP III promotion of cellular quiescence is required for the transition of host cells to syncytial cells, either for initial syncytial cell establishment or syncytial cell incorporation. These quiescent cells with partial xylem identity probably serve as a transitional stage for the host cell to fully transform into a syncytial cell. In support of this hypothesis, Anjam et al (2020) observed that BCN appeared to always select cells next to the xylem for syncytium initiation (Anjam et al., 2020), of which xylem adjacent (pro)cambial cells serves as stem cell organizer (Smetana et al., 2019), and xylem pole pericycle cells are competent for lateral root initiation with high auxin levels (Du and Scheres, 2018). Moreover, it has been reported that undifferentiated xylem precursor cells were always incorporated into the syncytium, buy not the fully differentiated xylem cells (Golinowski et al., 1996; Sobczak et al., 1997). In our observation, we also noticed that when the syncytium is formed near the root tip, xylem differentiation was often disrupted (Figure 8A; Supplemental Figure S16A). All these observations point towards the idea that xylem-identity cells could be an important transitional stage for syncytium formation. After the cell enters this transitional stage, other nematode effectors, or factors from the host cell, will then be needed to inhibit further xylem differentiation, and divert the cell into a syncytium. At this point, high auxin or HD-ZIP III expression would no longer be needed, and thus we saw down-regulation of *DR5::4xYFP* and *ProATHB8::4xYFP* expression in well-formed syncytium (Figure 9A-C, Supplemental Figure S17A-C). Interestingly, ectopic protoxylem cells were often seen at the edge of developed syncytia (Figure 9A-C), probably a result of insufficient signal in suppressing xylem differentiation.

The ectopic expression of *ATHB8d* in vascular cambium was sufficient to induce ectopic stem cell organizers, and to increase the number of xylem vessels, supposedly a result of shifted or increased stem cell organizer domains (Smetana et al., 2019). Nevertheless, moderate overexpression of *ATHB8d-YFP* with the *ANT* and *35S* promoters (Figure 7B) was not sufficient to increase BCN infection rate or syncytial cell size (Figure 7D-E). It is possible that in such a scenario, factors that promote transition of the stem cell organizer to a syncytial cell become limiting elements for syncytium establishment or expansion. Strikingly, high expression of *ATHB8d-YFP* driven by *G1090* promoter also suppressed BCN infection (Figure 7B, D). Such an extremely high level of ATHB8d expression may have tipped the balance between stem cell organizer specification and its further differentiation, thus preventing it from transiting into a syncytial cell.

Overall, in this study, we identified *ATHB8* as a downstream factor of CLE signaling in BCN infection in *Arabidopsis*, and showed that *ATHB8*, along with other HD-ZIP III factors, play important roles in syncytium cell formation, probably by promoting host cells into a quiescent status. However, the exact role of each individual member of HD-ZIP III in BCN infection, and their connection with CLE and auxin signaling would need further investigation.

### Data Availability

Raw sequence data have been submitted to the National Center for Biotechnology Information Sequence Read Archive under BioProject number PRJNA1054488.

## Acknowledgments

Special thanks to Clinton Meinhardt, Amanda Howland, Ben Averitt, Dean Kemp, and Kurk Lance for nematode population maintenance throughout the span of this project. We thank Dr. Nathan Bivens from University of Missouri DNA Core for his assistance in RNA seq library preparation. We thank Dr. Bill Spollen from University of Missouri Informatics Research Core for his assistance with RNA seq data analysis. We thank Dr. Muthugapatti K. Kandasamy of UGA Biomedical Microscopy Core for his confocal microscopy expertise and assistance. We thank Drs. Ari Pekka Mähönen and Annelie Carlsbecker for providing seeds.

## Author Contributions

X.L. and M.G.M. designed experiments; X.L. performed all experiments, analyzed the data, and drafted the manuscript; X.L. and M.G.M. revised the manuscript.

## Materials and Methods

### Plant materials

Plants are in Col-0 background unless otherwise specified. *clv1-101 clv2-101 rpk2-5* (Replogle et al., 2012), *ProATHB8::GUS* (Ws) (Baima et al., 1995), *ProATHB8::4xYFP* (Marques-Bueno et al., 2016), *Pro35S::XVE>>GUS*, *ProG1090::XVE>>GUS* (Siligato et al., 2016), *Pro35S::XVE>>miR165a*, *ProANT::XVE>>ATHB8d-YFP*, *ProMIR165a::GFP*, *ProMIR166b::GFP*, *MIR165/6-GFP* sensor line (Carlsbecker et al., 2010), and *ProCRE1::XVE>>MIR165a* (Muller et al., 2016), *ath8-11* (Col er-2 background), *cna-2* (Col er-2 background), *athb8-11 cna-2* (Prigge et al., 2005) have been described previously.

### Beet cyst nematode infection assay

*H. schachtii* was propagated on sugar beet (*Beta vulgaris* cv. Monohi). Nematode eggs were isolated and hatched as previously described (Mitchum et al., 2004). After 2 - 3 days, J2s (2^nd^ stage Juveniles) were collected and surface-sterilized with 0.004% Mercuric Chloride, 0.004% Sodium Azide and 0.002% Triton X-100 solution for 7 min, washed with sterilized water for 6 times, and then resuspended in 0.05% agarose. *Arabidopsis* seeds were sterilized and grown on modified Knop’s medium with Daishin agar (Brunschwig Chemie, Amsterdam, The Netherlands) (Sijmons et al., 1991).

For phenotypic analysis, seeds of each genotype were placed in 12 well plates following a random block design, stratified at 4 degree for 2-3 days, and were grown in a chamber set at 24 °C with 12 h light/12 h dark light cycle. Fourteen days after germination, 180-200 J2s were inoculated on each root. The penetration rate was counted 3 – 5 days after inoculation. Number of J4s/mature females were counted on 14th day and 30th day after inoculation. For syncytia size measurement, syncytia associated with single female were imaged on 15th day after inoculation and syncytia sizes were measured with ImageJ. When Estradiol induction is needed, 10 mM Estradiol stock in DMSO was first diluted to 25x of the working concentration in sterile water pre-warmed to 60 °C, and 100 ul of the diluted Estradiol or DMSO were added to each well (containing 2.5 ml Knop’s medium). Estradiol induction starts 2 days before inoculation.

### Penetration assay

Penetration rate was counted at 3 – 5 days post inoculation. Shoots of seedlings were first removed by scissors. The solid Knop’s media was melted by placing 12-well plates in a boiling water bath for 8-10 min and subsequently removed. Each well was rinsed with water twice to remove media residues. Nematodes were stained with boiling acid fuchsin solution for 1-2 min, rinsed with 95% ethanol once, and then kept in 95% ethanol till counting.

### Collection of infected root segments

To collect infected root segments, seeds were plated on square plates and grown vertically. About 10-15 sterilized *H. schachtii* J2s were inoculated on each root on the 7th day after germination, and root segments with syncytium (about 8 mm in length), or corresponding root segments on uninfected or HsCLE2p treated seedling roots, were cut under a stereoscope with scissors 4 days after inoculation. Root segments were immediately frozen in liquid nitrogen and were kept in -80 °C freezer until RNA isolation.

### RNA isolation

RNA was isolated by NucleoSpin RNA Plant Kit (Macherey-Nagel 1806/001) according to manufacturer’s instruction. RNA integrity was inspected by a bioanalyzer.

### Library construction and sequencing

RNA-seq library construction and sequencing were carried out in the University of Missouri DNA Core Facility. Libraries were constructed following the manufacturer’s protocol with reagents supplied in Illumina’s TruSeq mRNA stranded sample preparation kit. Briefly, sample concentration was determined by Qubit flourometer (Invitrogen) using the Qubit HS RNA assay kit, and the RNA integrity was checked using the Fragment Analyzer automated electrophoresis system. mRNA samples were enriched by poly-A enrichment and fragmented. Double-stranded cDNA was generated from fragmented RNA, and the index containing adapters were ligated to the ends. Amplified cDNA constructs were purified by addition of Axyprep Mag PCR Clean-up beads. Final construct of each purified library was evaluated using the Fragment Analyzer, quantified with the Qubit fluorometer using the Qubit HS dsDNA assay kit, and diluted according to Illumina’s standard sequencing protocol for sequencing on the NextSeq 500.

### RNA seq data analysis

For all samples, adapter sequence was removed using cutadapt (v0.16) (Martin, 2011). Reads were then mapped to *Arabidopsis thaliana* transcripts (Ensembl release 42) using Salmon (v0.12.0) (Patro et al., 2017). Transcript counts were converted into gene counts using the Bioconductor package tximport (v1.10.1) (Soneson et al., 2016). Differential expression of genes was tested using the Bioconductor package DESeq2 (v1.22.2) (Love et al., 2014). Gene ontology enrichment was analyzed by Bioconductor package topGO (v2.34.0) (Alexa and Rahnenfuhrer, 2018) with *Arabidopsis* annotation package org.At.tair.db (v3.7.0) (Carlson, 2018).

### qPCR

For qPCR validation, the same RNA samples for RNAseq were reverse transcribed using PrimeScript 1st strand cDNA Synthesis Kit (Cat# 6110A) according to the manual. qPCR was conducted with Applied Biosystem PowerUp SYBR Green master mix (cat# A25741) on a Bio-Rad CFX Connect Real-Time PCR System. 0.1 ul cDNA was used for each qPCR reaction.

Because typical references genes including *ACT2*, *ACT8*, *GAPDH, GAPB, UBP22,* and *UBC21* showed differential gene expression in our RNA sequencing dataset, a new set of reference genes were tested and *ACO3* (AT2G05710) was selected as the reference gene in our experiment (Supplemental Figure S5B-C). Primers used for qPCR were listed in Supplemental Table S2.

### GUS staining

For GUS staining of infection sites, seedlings were grown vertically on Knop’s medium, and inoculated with about 10-15 sterilized *H. schachtii* J2s 5-7 days after germination. Infected roots were collected on the indicated date and stained with GUS staining solution (1mM K Ferricyanide, 1mM X-Gluc, pH 7.0) overnight at 37 °C.

### Confocal imaging

Infected roots were collected at indicated time points and fixed in 4% paraformaldehyde (PFA) at 4 °C overnight, washed twice with 1xPBS, and were then cleared with ClearSee solution (Kurihara et al., 2015) supplemented with 0.5 ug/ml Calcofluor White (Sigma, Cat# 18909-100ml) for at least one week before imaging.

For live imaging, reporter lines were grown on ½ MS medium for 4 days, then were transferred to Knop’s media in a vertical plate. Each seedling was inoculated with about 10 sterilized *H. schachtii* J2s. About 18 hours after inoculation, seedlings that have been infected were selected under microscope and were moved to a block of Knop’s media (with 100 ug/ml Timentin) on a microscope slide, and then were covered with a 22 x 40 coverslip. The coverslip was secured by tape on both ends to prevent it from moving. The assembly was put in a square plate with the coverslip facing down and kept in a growth chamber (24 °C, 12/12 light/dark cycle). GFP/YFP signal was imaged on a Zeiss LSM 880 confocal microscope 1-5 days after inoculation, at approximately the same time each day.

## List of Supplemental Figures

Supplemental Figure S1. Expression of the *WOX4* gene upon HsCLE2p treatment.

Supplemental Figure S2. Quality control of RNAseq dataset.

Supplemental Figure S3. Overlapping genes down-regulated by BCN infection and HsCLE2p treatment in wild-type and the *clv* triple mutant.

Supplemental Figure S4. BCN infection reduced gene expression differences between *clv* triple mutant and wild-type roots.

Supplemental Figure S5. Selection of a new set of qPCR reference genes.

Supplemental Figure S6. qPCR verification of selected genes that are positively regulated by CLE signaling in BCN infection.

Supplemental Figure S7. Filtering for downstream genes that are negatively regulated by CLE signaling.

Supplemental Figure S8. Expression of HD-ZIP III transcription factors in the RNA sequencing dataset

Supplemental Figure S9. Expression of *ProATHB8::GUS* in 5-day old seedling of uninfected wild-type and the *clv* triple mutant.

**Supplementary Figure S10.** Expression of *ProATHB8::4xYFP* in uninfected root and BCN infection site (5 dpi)

Supplemental Figure S11. Expression of *ProMIR165a::GFP* and *ProMIR166b::GFP* in *Arabidopsis* root.

Supplemental Figure S12. Expression of *MIR165/166* regulatory genes, *AGO1*, *AGO10*, *SHR*, and *SCR* in the RNAseq dataset.

Supplemental Figure S13. Expression of *MIR165/6* sensor in BCN infection site in *Arabidopsis* root.

Supplemental Figure S14. Infection of *ATHB8d-YFP* overexpression lines in the vertical plates.

Supplemental Figure S15. Above-ground phenotype of *ATHB8d-YFP* overexpression plants in 12-well plates.

Supplemental Figure S16. Expression of auxin reporter *DR5::4xYFP* in the BCN induced syncytium at 1 dpi.

Supplemental Figure S17. Distribution of *DR5::4xYFP* activity at the BCN infection site at 3 dpi.

## List of Supplemental Tables

Supplemental Table S1. Mapping efficiency of RNA sequencing reads.

Supplemental Table S2. Primers used for qPCR.

## List of Supplemental Dataset

Supplemental Dataset S1. RNA-seq count table.

Supplemental Dataset S2. RNA sequencing differential gene expression analysis – all comparisons.

Supplemental Dataset S3. Gene Ontology (Biological Process) analysis of differential expressed genes.

Supplemental Dataset S4. List of genes positively regulated by CLE signaling in BCN infection.

Supplemental Dataset S5. List of genes negatively regulated by CLE signaling in BCN infection.

## Abbreviations

CN: Cyst Nematode
BCN: Beet Cyst Nematode
dpi: days-post-inoculation
J2: second stage juvenile

